# Collagen I is a critical organizer of scarring and CNS regeneration failure

**DOI:** 10.1101/2024.05.07.592424

**Authors:** Yihui Bi, Wenxiu Duan, Jerry Silver

## Abstract

Although axotomized neurons retain the ability to initiate the formation of growth cones and attempt to regenerate after spinal cord injury, the scar area formed as a result of the lesion in most adult mammals contains a variety of reactive cells that elaborate multiple extracellular matrix and enzyme components that are not suitable for regrowth^1,2^. Newly migrating axons in the vicinity of the scar utilize upregulated LAR family receptor protein tyrosine phosphatases, such as PTPσ, to associate with extracellular chondroitin sulphate proteoglycans (CSPGs), which have been discovered to tightly entrap the regrowing axon tip and transform it into a dystrophic non-growing endball. The scar is comprised of two compartments, one in the lesion penumbra, the glial scar, composed of reactive microglia, astrocytes and OPCs; and the other in the lesion epicenter, the fibrotic scar, which is made up of fibroblasts, pericytes, endothelial cells and inflammatory cells. While the fibrotic scar is known to be strongly inhibitory, even more so than the glial scar, the molecular determinants that curtail axon elongation through the injury core are largely uncharacterized. Here, we show that one sole member of the entire family of collagens, collagen I, creates an especially potent inducer of endball formation and regeneration failure. The inhibitory signaling is mediated by mechanosensitive ion channels and RhoA activation. Staggered systemic administration of two blood-brain barrier permeable-FDA approved drugs, aspirin and pirfenidone, reduced fibroblast incursion into the complete lesion and dramatically decreased collagen I, as well as CSPG deposition which were accompanied by axonal growth and considerable functional recovery. The anatomical substrate for robust axonal regeneration was provided by laminin producing GFAP^+^ and NG2^+^ bridging cells that spanned the wound. Our results reveal a collagen I-mechanotransduction axis that regulates axonal regrowth in spinal cord injury and raise a promising strategy for rapid clinical application.

## Introduction

Nerve repair is exceptionally difficult in the central nervous system (CNS), such as after spinal cord injury (SCI). CNS myelin-related growth inhibitors and CSPGs have been proposed to be the two principal extrinsic causes for regeneration failure. Methods to eliminate or overcome these inhibitors have stimulated several large-scale clinical trials^3–13^. However, neither enzymatic digestion nor genetic deletion nor receptor blockade of these growth barriers appear to promote frank long-distance axon regeneration across the fibrotic scar^10,12,14–16^. Thus, the removal of extracellular inhibitory molecules can promote mostly axonal sprouting^13,16,17^. In neurons with suppressed PTEN or activated mTOR^18,19^ that acquire strong intrinsic regenerative capacity, regeneration can occur directly across the lesion, but only if the lesion gap is quite narrow (<0.5 mm)^20^ allowing for cellular bridging through the otherwise forbidden fibroblast landscape^21^. These findings, coupled with the result that diminishing fibroblastic cells can promote modest lesion-crossing regeneration^21^, have highlighted the fact that the fibrous scar forms an exceptionally inhibitory microenvironment that is abundant with a matrix rich substrate. However, what happens to axons after encountering this boundary region is poorly understood. The previous perspective held that the fibrotic scar is a physical barrier mediated largely by collagen IV and its association with abundant basement membrane constituents that are mechanically obstructive as well as serving as a protein scaffold for other axon growth inhibitors^22–24^. Now, we have discovered that one particular matrix element in fibrotic scars, which is also the most abundant protein found in all vertebrates, collagen I, is a powerful mediator of dystrophic endball formation which, in turn, leads to exceptionally potent inhibition of axonal regrowth.

## Results

### Collagen I restricts axon outgrowth

Our laboratory has designed an intracellular sigma peptide (ISP) to modulate protein tyrosine phosphatase σ (PTPσ) and release CSPG-mediated axonal regeneration/sprouting inhibition after incomplete spinal cord injury (SCI)^12,25,26^. However, in the model of complete SCI caused by forceps compression, we have now learned that ISP rarely promotes the recovery of hindlimb motor function in both rats and mice, aligning with the outcome of a recent chondroitinase ABC (chABC) enzyme study^27^ (Fig 1a, c). We examined the characteristics of such severe lesion sites after ISP treatment. At 63 days post injury (dpi), immunostaining revealed that serotonergic (5-HT) axons that could navigate deeply within the glial scar, never-the-less, showed extensive endball formation and consistently stopped in the outermost edge of the fibrotic scar, identified in these experiments by collagen type I deposition (Fig 1b, d). Similar to other independent studies, 5-HT axons^1,12,16^, corticospinal tract (CST) axons^1,14–16,19–21^, propriospinal axons^28,29^ and dorsal root ganglion (DRG) axons^30^ all face significant challenges in traversing the scar center due to the presence of destructive activated macrophages^31,32^ and possibly its deposition of a dense membranous associated extracellular matrix (ECM) comprised of HS/CSPGs, laminin/nidogen, fibronectin and various collagens^33^. Previous studies have demonstrated dystrophic endball formation and a distinct over-adhered morphology of adult DRG axons in CSPG “coffee ring” spot gradients mixed with laminin but we^12,34^, as well as others^35^, have not investigated the effect of collagens in this 2-dimensional model. We now unveil that collagen type I is an overlooked strong inhibitory molecule for axon regeneration.

**Fig. 1.**
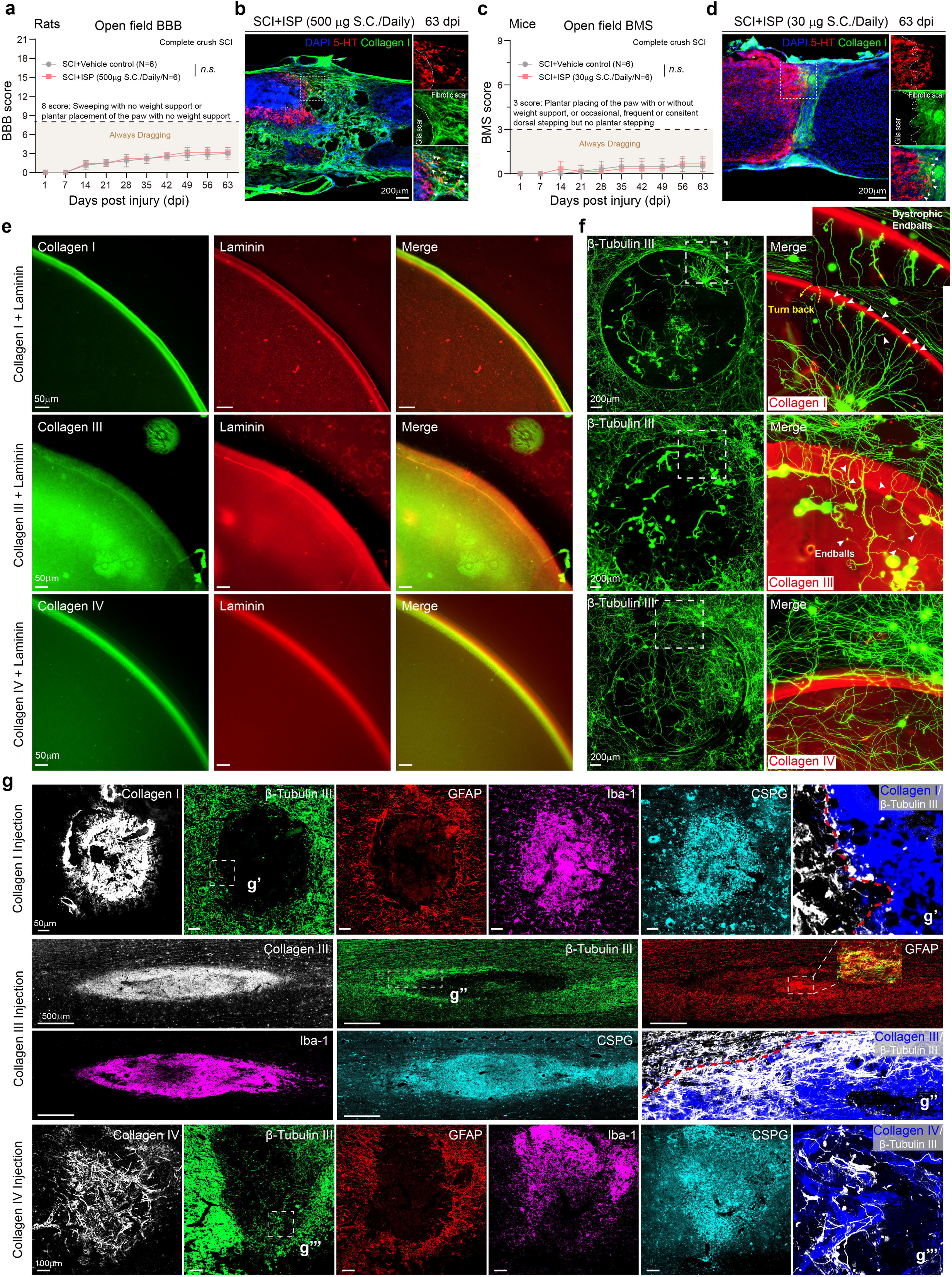
Identification of collagen I as an axonal growth inhibitor. **a**, **c**, Locomotor recovery after complete SCI (BBB score for rats and BMS score for mice). Error bars correspond to 6 animals per group. *n.s.*, not significant, two-way ANOVA test followed by post-hoc Bonferroni correction. The data are mean ± s.e.m. **b**, **d**, Images of spinal sections 63 days post injury (dpi) with antibodies against 5-HT and collagen I. DAPI, cell nuclei. Scale bars, 200 μm. **e**, Images of the collagen I/III/IV-laminin spot showing the distribution of the collagen I/III/IV (green) and laminin (red). The concentration of collagen I/III/IV and laminin are lower in the center of the spot and higher in the rim. Scale bars, 50 μm. **f**, Immunostaining of β-tubulin III (green) in a high-density dissociated adult DRGs culture on the collagen I/III/IV-laminin spot. The white arrows indicate the appearance of large club endings on neurites, and the yellow arrow highlights a neurite extending toward the collagen I-laminin spot rim, ultimately opting to reverse its direction. Scale bars, 200 μm. **g**, Images of spinal sections 7 dpi after collagen I/III/IV injection in ventral horn of rat spinal cord at T10 level stained with antibodies against the indicated proteins; Neuronal cell bodies and axons (anti-β-tubulin III), astrocytes (anti-GFAP), microglia and microphages (anti-Iba-1), CSPG (anti-CS56). Regions of interest for g’, g’’, and g’’’ are enlarged. Scale bars, 50 μm (collagen I injection), 500 μm (collagen III injection), and 100 μm (collagen IV injection).

Firstly, to elucidate the expression profile of collagens in SCI, we conducted RNA-sequencing analysis on spinal cords of adult mice subjected to crush injuries and identified wide changes of collagen expression involving an increase in fibrillar collagens, fibril-associated collagens, and network-forming collagens, highlighting the complex remodeling processes that occur following SCI (Extended Data Fig. 1a and Supplementary Table 1). *Col I (col1a1, col1a2), Col III (col3a1)* and *Col VI (col6a1, col6a2, col6a3)* mRNAs were the most abundant fibrillar/fibril-associated collagen members and were expressed highly specific to fibroblasts^36^ (Extended Data Fig. 1a, b). As the main type of network-forming collagen, *Col IV (col4a1, col4a2)* mRNAs were present at high levels in fibroblasts, pericytes, and endothelial cells (Extended Data Fig. 1a, b). Immunostaining revealed deposition of collagens I, III, IV and VI in damage centers from 7 to 28 dpi in mice (Extended Data Fig. 2a). The patterns of collagen I, III and IV expressions in rats closely mirrored those observed in mice, but collagen VI was rarely detected (Extended Data Fig. 2b, c).

As 5-HT axons were observed to establish direct contact with collagen I and develop extensive dystrophic endballs *in vivo*, this led us to investigate the role of collagens in adult axon regrowth. We harvested and then introduced fully mature rat DRG neurons onto a spot gradient of collagen mixed with the growth-promoting molecule laminin. Air dried collagen I, III and IV spots generated a crude gradient, wherein the periphery of the spot exhibited a significantly higher concentration of collagen compared to the center (Fig. 1e and Extended Data Fig. 3a). However, collagen VI failed to form a stable gradient spot *in vitro* (Extended Data Fig. 3b). Long-term exposure to high concentrations of collagen I is detrimental to neuronal survival and axon extension (Fig. 1f and Extended Data Fig. 3a). Fibers from the laminin surround located outside the spot regularly turned to extend along the sharp interface of the collagen I but seldom entered in. Furthermore, the DRG neurons surviving within the spot and especially those that invade the gradient from its inner edge appeared to be entrapped, as their axons were unable to extend freely and often formed a fasciculated structure with very large bulbous swellings at the tip. A few axons in the rim can turn back but these also develop dystrophic endings (Fig. 1f). This morphology bears a striking resemblance to Ramon y Cajal’s descriptions of chronically injured cat spinal cords^37^. In addition, DRGs growing on collagen III containing spots showed a behavioral pattern somewhat akin to that of collagen I, but, importantly, collagen III was considerably less inhibitory than collagen I (Fig. 1f and Extended Data Fig. 3c). On the contrary, collagen IV did not display discernible inhibition, aligning with findings from a prior study^38^ (Fig. 1f). Consistently, restricted outgrowth also occurred at the ends of axons of CNS derived hippocampal neurons exposed to a collagen I gradient (Extended Data Fig. 3d).

A typical fibrotic scar structure that forms in the damaged center of spinal cord lesions develops a more heterogenous mixture of dense depositions of basal lamina components such as collagen I, fibronectin, CSPGs and laminin^1,29^. This conglomeration of ECM molecules is far more complex than the two-molecule spot mixture described above. To better mimic the actual, *in vivo*, scar structure *in vitro,* we mixed various combinations of these four matrix components in the air-dried spot. Interestingly, as the four components were introduced one by one in the order of laminin-aggrecan-fibronectin-collagen I, a strikingly insoluble fiber precipitation emerged within the initially clear mixture upon the addition of collagen I in the final step. Immunostaining indicates that the fiber structure contains all four components, inspiring us to hypothesize that this new, more complete, *in vitro* model may better mimic the formation of fibrous scar matrix in SCI lesions (Extended Data Fig. 3e). The absence of laminin does not affect the formation of these fibers. On the contrary, collagen I plays the pivotal role, acting as a scaffold for binding to aggrecan and fibronectin^39^, which contribute to the insoluble fiber formation (Extended Data Fig. 3e-g). Interestingly, DRG axons again became trapped on the supramolecular fiber substrate and formed multiple endballs (Extended Data Fig. 3h).

Both neurons and astrocytes face hindrance in their growth within areas of collagen deposition following SCI. In alignment with prior findings^40,41^, our *in vitro* model demonstrated that astrocytes could adhere to the spot and those that grew within the spot center exhibited a condensed, highly reactive state (Extended Data Fig. 4a). However, while a few could invade the rim, most of those that made contact with the outer border coming from all directions did not cross. To serve as a control, it is important to note that fibroblasts exhibit unrestricted growth throughout the collagen I gradient spot (Extended Data Fig. 4b).

To test whether collagen I repels axons and astrocytes *in vivo*, we evaluated the state of the neuropil after stereotactic injection of collagen I into the ventral horn at level T10 of the spinal cord. Unlike lipopolysaccharide and various growth or inflammatory factors tested before^42^, we noticed that the signals for β-tubulin III and GFAP were conspicuously absent in areas where collagen I was present, indicating few or no neurons and astrocytes existed. By contrast, an intense accumulation of Iba-1^+^ microglia/macrophages was observed within the vicinity of collagen I deposits, coinciding with a substantial production of CSPGs (Fig. 1g). While the precise cellular origin of these CSPG accumulations was unclear, their patterning in relation to Iba-1 staining suggested that the proteoglycan might, in part, originate from these cells or, more likely, be phagocytosed by them^43^. Iba-1^+^ microglia/macrophages was incapable of degrading the collagen I and CSPG matrix, at least at the 1-week timepoint. We observed a somewhat comparable phenomenon at 1 week in spinal cords injected with collagen III. The distinction is that collagen III diffuses into an oval shape within the spinal cord, and some neurons and reactive astroglial cells manage to survive within the collagen III region. Intriguingly, it seems that the surviving reactive glia may facilitate neuronal survival in their immediate environment, aligning with the idea that early post-lesion reactive astrocytes can play a supportive role in maintaining neuronal/axonal integrity^44^ (Fig. 1g). At the injection site of collagen IV for one week, astrocyte loss, significant immune response, and CSPG deposition were also observed. Notably, we detected a large number of β-tubulin III positive nerve fibers growing on collagen IV, which is consistent with our *in vitro* results (Fig. 1g). Together, these results suggest that collagen I (with the help of collagen III and IV) appear to be pivotal factors contributing to the initiation of a cascade of cellular responses leading to an intense inflammatory response, the stimulation of CSPG production, rampant axotomy/dendritomy and astrocyte emigration out of the lesion core which abandons the neuropil to form a reactive glial boundary around the lesion.

### Mechanical stress determines the restriction

It has been hypothesized that the formation of endballs at the tips of regenerating DRG axons is a manifestation of multiple changes that arise from the stimulation by excessively tight substrate adhesion that can lead to a local synaptic-like phenotype which, in turn, may play a pivotal role in curtailing extension of the axon^30,34,45^. Although the dystrophic neuron is still alive, it can persist in this non-elongating physiological state for an extended duration *in vivo* (even many decades^46^). There are a number of markers that indicate that a synaptic-like phenotype has developed within such abnormal axon terminals. One of them is the amyloid precursor protein (APP). While APP is thought to be a damage response protein in neurons and serves as a general marker for axonal injury and dystrophy^47–49^, APP also helps to guide synapse formation^50,51^. Indeed, strong APP immunoreactivity appeared within “dystrophic endballs” upon collagen I substrates (Fig. 2a). The aggregated APP did not undergo transition to amyloid beta (Aβ) (Extended Data Fig. 5a). Similar results were observed after SCI *in vivo* with accumulation of APP in endballs in close association with collagen I expression (Fig. 2b, Extended Data Fig. 6). We also observed a significant increase in the intensity of presynaptic markers, such as synaptic vesicle glycoprotein 2 (SV2) and vesicular glutamate transporter 1 (V-Glut1) immunostaining, within “dystrophic endballs” (Extended Data Fig. 5b, c). Tubulin associated unit (Tau-1) immunostaining revealed no major structural changes in axonal integrity (Extended Data Fig. 5c). Thus, our results suggest that collagen I restricts axonal migration by triggering growth cones to form an APP-filled synaptic-like structure that enters into a dystrophic, non-elongating condition.

**Fig. 2.**
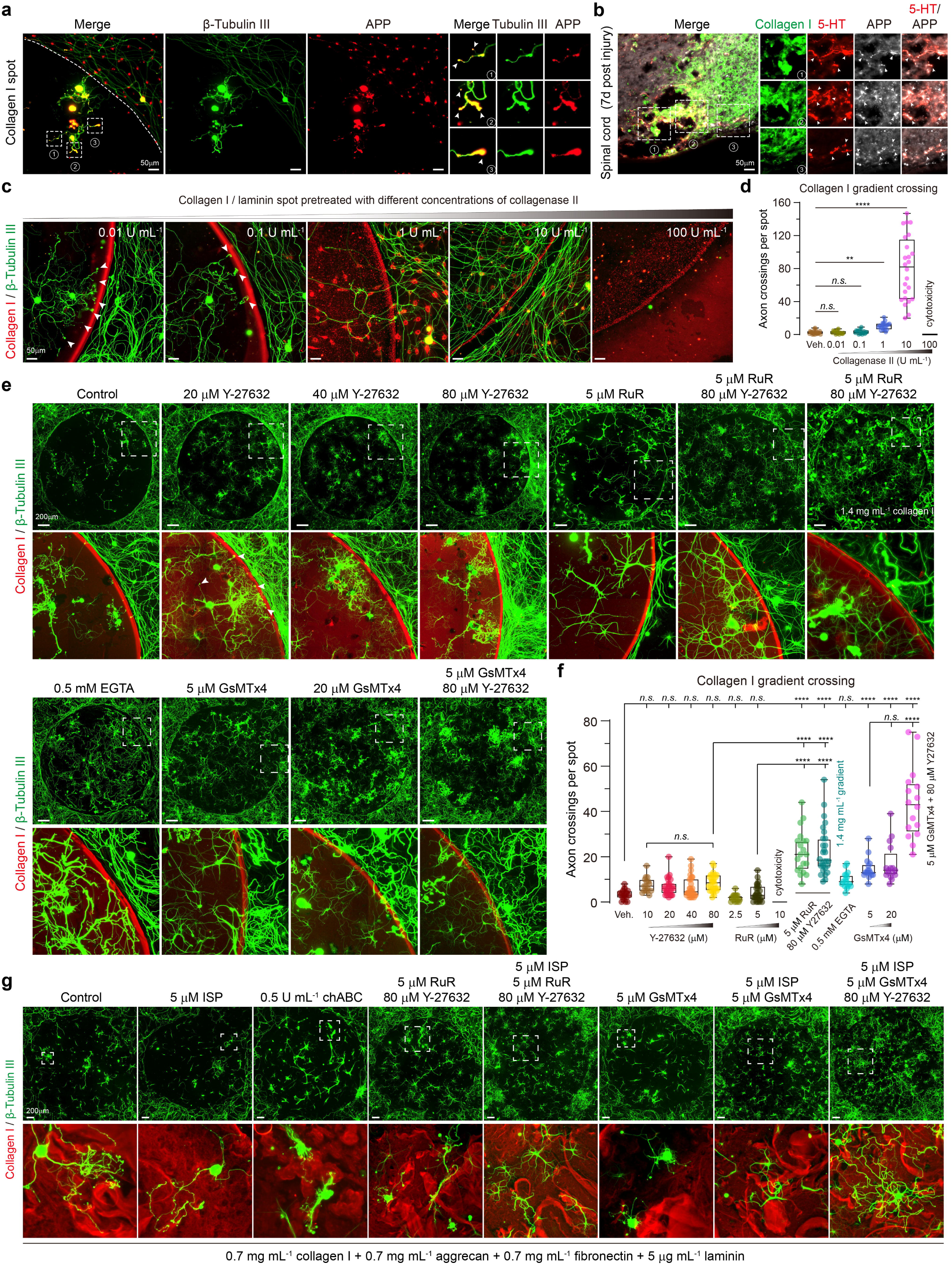
Collagen I acted as a biomechanical modulator rather than a signaling molecule. **a**, Representative images of “dystrophic endballs” immunolabelled for β-tubulin III and APP (Amyloid precursor protein). Regions of interest for 1, 2, and 3 are enlarged, indicating that the APP is enriched in endballs. **b**, Images of spinal cord sections in mice at 7 dpi with antibodies against collagen I (green), 5-HT (red), and APP (white). **c**, Representative images of DRGs growing on collagen I gradient spots pre-treated with collagenase II. The white arrows indicate the appearance of “dystrophic endballs”. **d**, Quantification of numbers for DRG axons crossing the inhibitory rim (*n* ≥ 16 per group). \*\**p* = 0.0022, \*\*\*\**p* < 0.0001, *n.s.,* not significant, Kruskal-Wallis test followed by post-hoc Dunn correction. Data are min to max, show all points. **e**, Microscopy images of β-tubulin III (green) in DRGs on a complete collagen I gradient spot (red) under a series of drug exposures. **f**, Quantification of numbers for DRG axons crossing the inhibitory rim (*n* ≥ 16 per group). \*\*\*\**p* < 0.0001, *n.s.,* not significant, one-way ANOVA followed by post-hoc Bonferroni correction. Data are min to max, show all points. **g**, Microscopy images of β-tubulin III (green) in DRGs on a complete collagen I (red)-aggrecan-fibronectin-laminin spot under a series of drug exposures. Representative regions of each group are magnified for display (**e**, **g**). Scale bars, 50 μm (**a**, **b**, **c**), 200 μm (**e**, **g**).

Next, we attempted to explore strategies to release collagen I-mediated axonal restriction *in vitro.* Gradually increasing the concentration of the growth-promoting molecule laminin to 25 μg/mL, which was shown to be effective in CSPG gradients^34^, surprisingly failed to allow axons to extend across the collagen I gradient (Extended Data Fig. 7a, b). Even though a statistically significant increase in the number of axons entering the rim was observed upon amplifying the laminin concentration all the way to 100 μg/mL, this was still insufficient to overcome the tendency to form dystrophic endballs. Pre-treatment with collagenase II, which breaks the triple-helical structure in fibrillar collagen^52^, promoted robust axonal regrowth through the collagen I gradient in a dose-dependent manner (Fig. 2c, d). Unfortunately, the most effective, growth permitting doses of collagenase became toxic to neuronal cells. Collagenase also disrupts the extracellular matrix of intact blood vessels when administered *in vivo*^53^, markedly diminishing its potential for therapeutic application. Additionally, neither ISP nor chABC as well as the microtubule-stabilizing drug ixabepilone (a stable, long acting epithilone) ameliorated the inhibitory activity of collagen I (Extended Data Fig. 7c, d).

Previous studies^54^ have shown that collagen I can act as a signaling molecule to regulate cell migration and adhesion through various collagen receptors, such as integrins (α1β1, α2β1, α10β1 and α11β1), discoidin domain receptors (DDR1 and DDR2), leukocyte receptor complexes (Ocsar, Gp6, and Lair1), and mannose receptors (Mrc1, Mrc2, Pla2r1 and Ly75) (Extended Data Fig. 8a). Single-cell sequencing data^55^ indicated that DRG neurons mainly express integrin-β1and the DDR1 receptor (Extended Data Fig. 8b). Moreover, neurons in the spinal cord^36^, cortical motor neurons^56^, and caudal raphe neurons^57^ all express these two receptors (Extended Data Fig. 8c-e and Extended Data Fig. 10a, b). We initially speculated that the inhibitory effect of collagen I might be mediated by one or both of these two receptors. Nevertheless, neither the function-blocking monoclonal antibody targeting integrin-β1 (anti-β1 Ab) nor DDR1 inhibitors, including nilotinib, merestinib, and sitravatinib were able to reverse the restriction (Extended Data Fig. 8f, g). Considering the prior evidence of collagen binding to G-protein coupled receptors (GPCRs)^58,59^, our attention shifted to G protein-coupled receptor 56 (GPR56), a pivotal GPCR involved in neuron development and neurite growth^60^. All the neurons we were focusing on exhibited GPR56 (*Adgrg1*) expression, except for cortical motor neurons (Extended Data Fig. 8h-j and Extended Data Fig. 10b). Therefore, we assessed the effects of recombinant PLL domain (rPLL-rbFc), known to inhibit GPR56 binding to its ligand, as well as dihydromunduletone (DHM), a GPR56 inhibitor. However, again, no significant changes were observed in our spot assay model (Extended Data Fig. 8k, l).

Next, we attended to accumulating evidence which suggests that substrate rigidity and mechanosensing play pivotal roles in regulating axonal growth and regeneration^61,62^, which led us to speculate that the collagen I matrix might act as a biomechanical modulator rather than a more molecular transduction form of signaling molecule. Notably, previous studies have indicated that early forming glial scars (lacking collagen I) are softer than surrounding tissue^63^. However, recent research using atomic force microscopy has discovered that fibrotic scars progressively stiffen as collagen I accumulates^64^. Since RhoA signaling is central to mechanotransduction^65^, we tested Y-27632, an effective RhoA inhibitor. Diminishing RhoA resulted in an increase in neuronal survival and a reduction in dystrophic endball formation (Fig. 2e). However, even after increasing the concentration of Y-27632 to a high level, most axons entering the rim still failed to cross freely (Fig. 2e, f). A number of mechanically gated ion channels are responsible for mechanotransduction and have been reported to contribute to the inhibition of axon regeneration^66^. As a non-selective mechanosensitive ion channel inhibitor, we next tested ruthenium red (RuR). Indeed, RuR alone also reduced somewhat the formation of endballs (Fig. 2e). We next tested the combination of both drugs. Surprisingly, even at high concentrations of collagen I, the combination treatment of RuR and Y-27632 allowed DRG neurons to extend axons readily through the gradient interface (Fig. 2e, f). EGTA promoted the survival and extension of axons within the spot, suggesting that these mechanically sensitive channels might be calcium selective. However, EGTA failed to improve axonal crossing significantly. Furthermore, GsMTx4, a more selective cationic mechanosensitive channel inhibitor, was also effective in our gradient model which provided deeper insight into the notion that collagen I-mediated restriction might be attributed to Piezo, TRP, BK and/or TACAN calcium channels^67,68^ (Fig. 2e, f).

We next turned to our new, 4 matrix molecule *in vitro* model of the fibrous scar to see what might be able to alleviate its powerful inhibition of axonal regrowth. Notably, we observed that solely overcoming CSPGs by blocking the PTPσ receptor with ISP or removing CS GAGs with chABC were insufficient to promote the extension of DRG axons on the more complex fibrous scar model substrate. However, a significant improvement in axonal extension capacity was achieved when the inhibitory effects of both CSPG and collagen I were simultaneously mitigated via ISP and mechanoreceptor modulation (Fig. 2g, far right panel). We next focused our attention *in vivo* to demonstrate that Piezo, TRP, BK and TACAN receptors are widely distributed in DRG and spinal cord neurons, cortical motor neurons, and caudal raphe neurons (Extended Data Fig. 9a-d and Extended Data Fig. 10a-b). Despite the absence of more specific pharmacological tools for tracking the contribution of each type of mechanoreceptor to the inhibitory effects of our model, our data and analysis still suggest that mechanical stress induced by collagen I is a major determinant in the formation of dystrophic growth cones and restriction on axonal growth.

### Combining aspirin with pirfenidone promotes scar-free repair, axonal regeneration and functional recovery after SCI

We next became dedicated to discovering a way to target the deposition of collagen I within the injured spinal cord via a clinically feasible and readily translational manner. Aspirin^69^ and pirfenidone^70,71^ are two FDA-approved orally deliverable drugs that have widespread clinical use, and both have demonstrated the ability to readily penetrate the CNS and alleviate fibrosis (Fig. 3a, b). Immunoblotting confirmed that both aspirin and pirfenidone reduced the production of collagens I and III in rat meningeal fibroblasts in a concentration-dependent manner *in vitro* (Fig. 3c, d and Extended Data Fig.11 and 12a). Given that the effective concentrations of the two drugs used individually are relatively high (5 mM each), and there are established daily intake dose limits in clinical settings, we combined these two drugs and observed a significant synergistic effect when the drug concentrations were raised to just 3 mM each (Fig. 3e, f and Extended Data Fig. 12b). In our collagen I gradient spot, aspirin and pirfenidone treatment (3 mM) has no obvious toxicity and also no effect on axon growth in or outside the outer rim interface (Extended Data Fig. 12c, d). Recall that in this assay collagen I is already present and aspirin/pirfenidone do not eliminate pre-formed collagen.

**Fig. 3.**
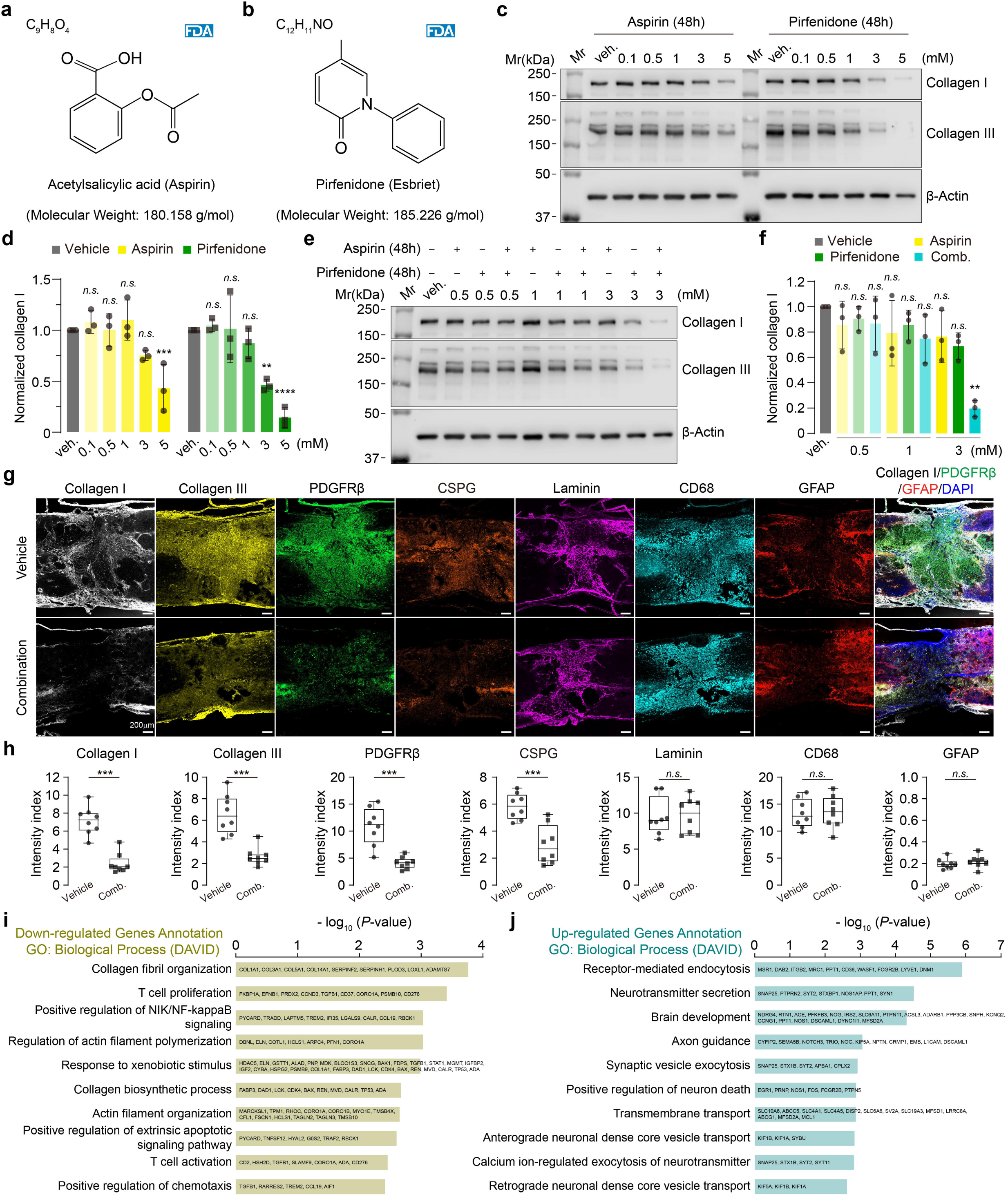
Aspirin and pirfenidone treatment promotes scar free wound healing in SCI. **a**, **b**, Chemical structure of aspirin (**a**) and pirfenidone (**b**). **c**, Primary rat meningeal fibroblasts treated with aspirin or pirfenidone alone were examined for collagen I and collagen III 48 h after treatment. **d**, Quantification of the indicated intensity (normalized to each β-actin and the vehicle control) in **b**. Data are mean ± s.d. *n* = 3, three independent tests. \*\**p* = 0.0014, \*\*\**p* = 0.0008, \*\*\*\**p* < 0.0001, *n.s.,* not significant, two-way RM ANOVA followed by post-hoc Dunnett correction. **e**, Expression of the collagen I and III in primary rat meningeal fibroblasts following aspirin and pirfenidone combination treatment. **f**, Quantification of the indicated intensity (normalized to each β-actin and the vehicle control) in **e**. Data are mean ± s.d. *n* = 3, three independent tests. \*\**p* = 0.0076, *n.s.,* not significant. Kruskal-Wallis test followed by post-hoc Dunn correction. **g**, Immunostaining of collagen I, collagen III, PDGFRβ, CSPG, laminin, CD68, and GFAP in injured core at 14 dpi after treatment. **h**, Quantification of immunoreactive intensity for **g** (normalized to the intact region). Data are min to max, show all points (n = 8). \*\*\**p* = 0.0003 (collagen I and collagen III, Mann Whitney test), \*\*\**p* = 0.0007 (PDGFRβ, Unpaired *t* test with Welch’s correction), \*\*\**p* = 0.0005 (CSPG, Unpaired *t* test Welch’s correction), \*\*\*\**p* < 0.0001 (collagen I), *n.s.,* not significant, Unpaired t test with Welch’s correction. **i**, **j**, Selected biological process terms and associated genes altered by aspirin and pirfenidone combination treatment at 14 dpi after SCI relative to vehicle treatment control (*n* = 3), as identified by Gene Ontology (GO) analysis on DAVIA (v2023q2). Scale bars, 200 μm (**g**).

To test whether the combination of aspirin and pirfenidone could ameliorate acutely forming fibrotic scar progression in adult rats following SCI, rats were administered aspirin (45 mg/kg BW, once daily, started at 8 dpi) and pirfenidone (16 mg/kg BW, three times a day, Days 1 through 7; 32 mg/kg BW, three times a day, Days 8 through 14; 48 mg/kg BW, three times a day, Days 15 onward until 56 dpi), following the already established clinical governance of the administration of these drugs (Extended Data Fig. 12e). At two weeks after treatment, the collagen I/III area and PDGFRβ^+^cell numbers were significantly decreased compared with vehicle controls (Fig. 3g, h). Notably, the lesions in rats in the aspirin and pirfenidone treatment group, exhibited a significant reduction in CSPG deposition, whereas, surprisingly, laminin levels remained unchanged (Fig. 3g, h). This reduction in CSPGs is likely an indirect effect of the treatment since it has been shown that collagen I is a potent inducer of glial CSPG production^40^. In addition, CD68^+^ cell accumulation, overall astrogliosis and lesion area were similar between treated and control rats (Fig. 3g, h).

To probe more broadly the influences of aspirin and pirfenidone treatment in SCI lesions, we conducted bulk RNA-sequencing of the injured spinal cord at two weeks after SCI. At this time point, aspirin and pirfenidone mainly downregulated genes related to collagen fiber organization and collagen biosynthetic processes (Fig. 3i). Notably, the upregulated genes were enriched in those related to neurotransmitter secretion, brain development, and axon guidance, which may suggest that fibrillar collagen-free induced lesions have specific molecular properties that allow or even promote axon regeneration (Fig. 3j).

Considering the promising clinical applications of aspirin and pirfenidone, we next asked whether the combined administration of these two drugs could facilitate functional recovery, specifically enhancing motor ability in the hindquarters, in rat models of SCI. We conducted tests on two types of incomplete injury models: incomplete crush and contusion SCI, and one type of complete injury model: complete crush SCI. We first performed horizontal ladder-walking tests^21^ in three separate cohorts of rats that underwent incomplete crush injury and received the combination treatment. Incomplete crush at T10 caused severe motor impairments immediately which lingered during the first week (Fig. 4a and Extended Data Fig. 13a-c). There was a gradual and continuing modest recovery that occurred over the next several weeks. In comparison to the vehicle control cohorts, significant recovery of function was observed starting from the third week post-treatment, characterized by much improved coordination of the trunk and fewer hind limb errors (Fig. 4a, b, Extended Data Fig. 13a-d, and supplementary movie 1). We focused further analysis on the serotonergic system, whose axons are relatively easy to visualize and contribute greatly to proper neuromodulatory tone in locomotion circuitry^12,72^. In rats with functional improvements after aspirin and pirfenidone treatment, we observed much higher numbers of 5-HT fibers in the lesion core and their enhanced presence well below the lesion (Fig. 4c and Extended Data Fig. 13e).

**Fig. 4.**
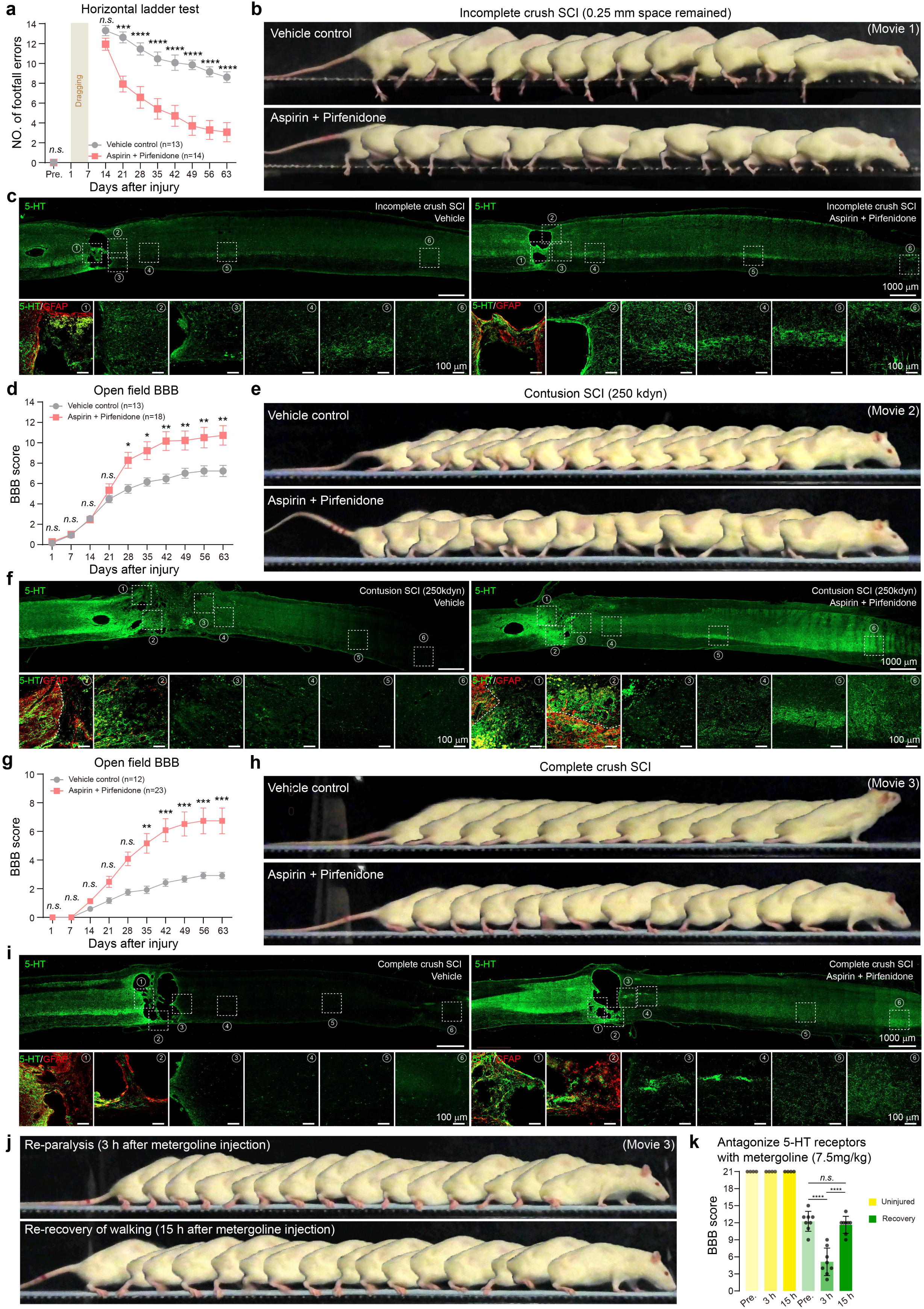
Functional recovery after Aspirin and pirfenidone combination treatment in SCI. **a**, Locomotor recovery was quantified using horizontal ladder-walking test in incomplete crush model (0.25 mm space remained) (*n* = 13 rats in vehicle group, *n* = 14 rats in aspirin and pirfenidone group). \*\*\**p* = 0.0002, \*\*\*\**p* < 0.0001, *n.s.*, not significant, two-way RM ANOVA followed by post-hoc Bonferroni correction. Data are mean ± s.e.m. Raw data are provided in **Extended Data Fig. 14a-d**. **b**, Chronophotography of walking in horizontal ladder-walking test at 9 weeks post SCI. **c**, 5-HT intensity in representative sagittal sections at 9 weeks post SCI showing serotonergic axons. The 5-HT fibers in injury lesion and caudal spinal cord are magnified. **d**, Locomotor recovery (BBB score) in severe contusion model (250 kdyn) (*n* = 13 rats in vehicle group, *n* = 18 rats in aspirin and pirfenidone group). \**p* < 0.05, \*\**p* < 0.01, *n.s.*, not significant, two-way RM ANOVA followed by post-hoc Bonferroni correction. Data are mean ± s.e.m. Raw data are provided in **Extended Data Fig. 14f-j**. **e**, Chronophotography of walking in contusion model at 9 weeks post SCI. **f**, Representative images from sagittal sections showing serotonergic axons in vehicle-treated rat and combination-treated responders at 9 weeks post SCI. **g**, Locomotor recovery (BBB score) in complete crush model (0.2 mm tip forcep) (*n* = 12 rats in vehicle group, *n* = 23 rats in aspirin and pirfenidone group). \*\**p* = 0.0024, \*\*\**p* < 0.001, *n.s.*, not significant, two-way RM ANOVA followed by post-hoc Bonferroni correction. Data are mean ± s.e.m. Raw data are provided in **Extended Data Fig. 14n-q**. **h**, Chronophotography of walking in complete crush model at 9 weeks post SCI. **i**, Representative images from sagittal sections showing regenerated serotonergic axons projection in rats with substantial walking recovery (BBB = 13) at 9 weeks post SCI. **j**, Chronophotography of walking in responders that received metergoline injection to antagonize the function of 5-HT. **k**, Walking was quantified using BBB rating score after metergoline injection (*n* = 4 uninjured rats, *n* = 8 responders). \*\*\*\**p* < 0.0001, *n.s.*, not significant, one-way ANOVA followed by post-hoc Bonferroni correction. Data are mean ± s.d.

We then turned to a severe (250 kdyn) contusive SCI at the T10 segment, and measured locomotor recovery using the Basso, Beattie and Bresnahan (BBB) 21-point rating scale^73^. After contusive SCI, vehicle-treated rats recovered slowly from initial hindlimb paralysis to weak spontaneous sweeping movements at week 8 (average BBB = 7.2) (Fig. 4d, e and Extended Data Fig. 13f-h). Aspirin and pirfenidone treatment resulted in a progressive recovery of leg movements that emerged ∼3 to 4 weeks after SCI. 39% (a total of 7 out of 18) rats displayed consistent weight support and forelimb-hindlimb coordinated stepping (BBB ≥ 14), with 2 rats achieving a gait pattern approaching that of uninjured rats (BBB = 18), and on average, the BBB score of all treated animals was 10.7 (Extended Data Fig. 13i, j and supplementary movie 2). Anatomical analyses confirmed that the cord contusions led to a remarkable decrease in descending 5-HT fibers caudal to the lesion (Fig. 4f). Instead, aspirin and pirfenidone treated responders showed robust 5-HT fibers that had either regrown across the glial/fibrotic scar or sprouted from spared axons leading to an enhanced serotonergic axon accumulation in the lumbar spinal cord (Fig. 4f and Extended Data Fig. 13k).

Incomplete crush and contusion SCI allow for consistent lesion bridging that leads to highly variable white matter sparing^12,72^, making it elusive to understand whether lengthy regeneration across and beyond the lesion might result in an abundance of 5-HT fibers in the lower spinal cord. The anatomically complete SCI model eliminates the complication of deciphering the role of distal axonal sprouting^19,28,29^. Rats that underwent complete crush SCI with a wide-tipped forceps (0.5 mm width) exhibited consistent paralysis, even at 8 weeks after injury (Extended Data Fig. 13l). Nevertheless, continuous monitoring of behavior over an 8-week period did not reveal improvements in the 5 aggressively crushed rats treated with aspirin and pirfenidone. Anatomical analyses found that except for one rat in which collagen was not significantly inhibited in the spinal cord, the remaining four rats had developed extensive cavities at the injury lesion (1.36 ± 0.11 mm), as previously reported^21,74^ (Extended Data Fig. 13l, m). Such rats exhibited little or no 5-HT axons with the ability to navigate the lesion. We reasoned that complete, wide diameter crush lesions led to an excessively large area of tissue damage and lack of potential cellular bridging to support regeneration across the core, even after fibroblast proliferation and collagen I/III synthesis were inhibited.

It is reasonable to speculate that any spontaneous axonal regeneration across a complete lesion requires some sort of a cellular bridge^19,20,29^, lacking densely packed fibrotic components^1,21,30^. We posited that creating a narrower lesion may help preserve the tissue bridges needed for regeneration. To test this possibility, we engineered a very fine forceps (0.2 mm width) to replace the standard one (0.5 mm width). We observed in three independent experiments of 23 rats that a total of 8 treated rats regained variable degrees of hindlimb walking (baseline average BBB =2.9 to average BBB = 6.7), with some attaining rather remarkable improvements (BBB = 9-15, average score of responders = 12) (Fig. 4g, h and Extended Data Fig. 13n-q). However, in the ladder test, the hindlimbs of lesion-complete rats with overground walking recovery still made many foot slips through the ladder rungs (Extended Data Fig. 14). In vehicle treated rats, if the glial and fibrotic scar borders occurred within a tissue bridge, then 5-HT axons that could regenerate as far as this interface, none-the less, stopped abruptly at the position of strong fibrillar collagen I deposition (Extended Data Fig. 15a). Aspirin and pirfenidone treatment facilitated robust 5-HT axon regrowth in responders (Fig. 4i and Extended Data Fig. 13r). In these rats, 5-HT fibers passed through newly forming reactive astrocyte borders, continued along bridges of reactive astroglial as well as non-astroglial cells across the lesion core and upon exiting the lesion that were lacking in fibrillar collagen I expression and upon exiting the lesion, projected well into the gray matter reaching as far as the lumbar spinal cord (Extended Data Fig. 15a, c). By interrogating a potential molecular mechanism of regrowth, we observed that regenerating 5-HT axons elongated on greatly reduced collagen I, excess laminin producing cell bridges that spanned the lesion core and they avoided CD68^+^macrophages^20,29,75,76^ (Extended Data Fig. 15a-d). Many of the laminin^+^ cells were closely associated with blood vessels especially in the tissues adjacent to the doorways of the bridge, suggesting that these cells are likely to be pericytes (Extended Data Fig. 15d arrows). The regenerated axons extended along both GFAP^+^ astrocytes as well as cells that expressed the NG2 chondroitin sulfate proteoglycan^20,29,43,77,78^ (Extended Data Fig. 15e). It was readily apparent that most regrowing axons in the lesion core associated most intimately with NG2^+^ cells (Extended Data Fig. 15d, f). It is also important to note that the astrocyte scar border cells in association with the re-growing axons aligned their shapes to orient more linearly with the axons (Extended Data Fig. 15e). Thus, our present results suggest that tissue bridges lacking fibrillar collagens can facilitate reactive astroglial plasticity^40^ which helps in the creation of a permissive environment across a lesion, at least for adult serotonergic axon regrowth after an acute complete SCI. However, we must also stress mention that over 65% of rats with narrower forceps lesions still did not respond to treatment. Anatomical analyses of these non-responding rats revealed large cavernous voids without tissue bridges emphasizing that our current strategy cannot control the assembly of residual wound spanning tissue.

Finally, we tested the necessity of serotonergic projections in the recovery of hindlimb walking after complete SCI. Pharmacological inhibition of serotonergic innervation by metergoline impaired walking in all responders at 3 hours after injection. (Fig. 4j, k and supplementary movie 3). After 15 hours, when metergoline was metabolized, these rats regained their walking ability. By contrast, metergoline administration in intact rats did not induce detectable deficits of locomotion (Fig. 4k). These results established that long-distance regeneration of serotonergic projections contributes to the substantial recovery of walking after complete SCI. However, serotonergic projections are unlikely to be responsible for total control of hindlimb behavior independently, suggesting that they are likely enhancing the functional regrowth of other descending tracts essential to restoration of movements, such as the corticospinal, propriospinal or reticulospinal systems^28,72^. We are now examining new cohorts of complete crush animals to determine which other supraspinal systems may be regenerating.

## Discussion

The phenomenon of fibrosis is one of the most devastating, life threatening sequelae of inflammatory processes that can occur in nearly every organ system in the body, including the CNS^71,79^. While the fibrotic scar has long been recognized to constitute a highly repulsive barrier for spontaneous axon regrowth in adults after SCI^80–82^, it has recently garnered more intense interest after it was shown that regeneration across the lesion was increased following the reduction of fibrotic cells^21,79,83^. Given the critical role of fibrosis in regeneration failure, it is important to understand the more precise molecular mechanisms underlying how fibrotic scar organizes this regrowth failure. We anticipated that this strong inhibition may be coordinated by one or a cluster of unsupportive/non-permissive extracellular substrates. Of the four well known components involved in fibrotic scar formation, the growth promoting properties of laminin and fibronectin^1,29,34^, and the inhibitory functions of CSPGs have been widely established^9–13,17,29^. Remarkably, the regeneration inhibiting potential of collagens has been largely neglected beyond the compendium of work beginning in the late 1990’s from the lab of Hans Werner Müller, who focused mostly on the meshwork of basal laminae and associated collagen type IV that develops after lesions of the adult CNS^22,82,84^. With the use of a variety of collagen IV inhibitors, the Muller lab purported to reduce basal lamina and promote functional regeneration of several different CNS axon tracts in the brain and spinal cord^84^. However, the lingering dilemmas around this hypothesis were, firstly, that Cajal believed the growth cone to be a “battering ram” capable of piercing its way through basement membranes (e.g., during development of the 8th cranial nerve as it emanates from the vestibulocochlear ganglion and invades through the basal lamina to enter the hindbrain^85,86^) and, secondly, that collagens in general and particularly collagen IV per se are not inhibitory for axonal outgrowth^22,23^. So, it was postulated that collagen IV served as a binding matrix for numerous other extracellular components and through its close interactions with known inhibitory molecules like proteoglycans and semaphorins blocked regeneration^87^. Thus, the overtly inhibitory properties of collagen I came as a surprise, especially since collagen I has long been used to create scaffolds for cell and axon growth in the CNS^88^.

To address this question, we first characterized the capacity of collagen I to restrict DRG axon outgrowth by causing the formation of remarkably large dystrophic endballs, unlike those seen before in our aggrecan/laminin spot assay model *in vitro*^34^. Interestingly, such massive endings have recently been described on the central processes of crushed sensory axons that can regenerate in the root but fail to enter the spinal cord where they abut collagen I bearing tissue at the dorsal root entry zone (DREZ)^30^. A similar type of ending, but one that is much smaller and even more synaptic-like, becomes entrapped at the DREZ as well as in the glial scar penumbra of an SCI in a tight association with NG2 chondroitin sulfate proteoglycan producing oligodendrocyte progenitor cells, where it can remain for many years^30,45^. Whether the large dystrophic endballs can remain permanently fixed in association with collagen I producing cells is unknown and will need to be studied in more chronically surviving lesion models. The intracellular mechanism by which CSPGs convert the growing terminal of the axon into an endball state is thought to be mediated, at least in part, via disruption of the process of autophagic flux linked to PTPσ dephosphorylating interactions with cortactin, a major actin binding protein involved with autophagosome fusion with lysosomes^35^. In contrast to such traditional molecular transduction forms of inhibitory signaling we have identified that the rigidity of the collagen I substrate likely acting through various mechanoreceptors determines the restriction. However, whether mechanoreceptor interactions with collagen I also instruct endball formation by disrupting autophagy still needs to be investigated. While we have not yet identified the specific mechanosensitive ion channel that mediates the inhibitory effect of collagen I, these channels seem to have different expression patterns in different descending and ascending neurons. Exploring these channels on functionally relevant neuronal subpopulations will be interesting, as strategies targeting these channels are likely to extend to the treatment of chronic SCI.

Two additional questions still need to be resolved: what is the precise cellular source of lesion core associated collagen I? what is the sequence of biological events that forms the regeneration bridge across the lesion? While it is abundantly clear that fibroblasts produce collagen I (*col1a1* and *col1a2*), and although identifying pericytes remains a challenge, the prevailing view is that pericytes do not express collagen, especially collagen I^89,90^. In mice, A-type pericytes identified or defined by Glast-cre (*Slc1a3*) were proposed to be the predominant cell type that forms the lesion core of fibroblast-like cells and dense extracellular matrix in SCI^21,74^. Although Glast-cre was often used to target glia and neural progenitor cells, fibroblasts also express *slc1a3* in SCI^36,91^. Thus, it remains debatable whether A-type pericytes in the CNS can become or are already fibroblastic^92^. Independent studies have also confirmed that actual fibroblasts, which envelop the large-diameter vessels, rather than pericytes which surround small caliber blood vessels, promote the formation of collagen I encumbered fibrous scars after CNS damage^83,93^. NG2^+^ cells (oligodendrocyte precursor cells and/or pericytes) were identified to curtail axonal regeneration by expressing NG2 (CSPG4) or inducing synaptic-like connections^30,45,76^. This proteoglycan mediated inhibition was considered as relative rather than absolute, especially when increases in laminin occur or CSPGs are cleared^29,34,76^. Thus, multiple changes occur following our combined drug treatment to the collagen I laden, lesion spanning cellular bridges which are comprised of several cell types including astroglia, fibroblasts, pericytes, macrophages and likely also endothelial cells. After treatment, fibroblasts are inhibited and collagen I is reduced, lesion CSPG deposition decreases likely also due to the lack of fibroblast produced collagen I, the morphology of glial scar border cells in the presence of regenerating axons is realigned which allows axons to migrate further into the bridge to interact with enhanced pericyte laminin which facilitates axon crossing^76–78^.

Even though we still need to mend the gap to create more bridges for axon regrowth, inhibition of fibrillar collagen (especially collagen I) after SCI in addition to enhancing the pericyte’s scaffold forming abilities should be applied as a fundamental framework in the testing of further combinatorial therapies, as this strategy of relieving fibrous scar inhibition along potential paths across the lesion may have a substantial promoting effect on regeneration leading to better functional recovery.

## Methods

### In vitro assay

#### Primary rat DRG and satellite glia culture

DRGs were obtained from adult female Sprague-Dawley rats (170-190 g weight) as previously described^12,34^. First, the DRGs were removed from the waist to the neck on ice and the cell bodies were carefully dissected out under a microscope. The DRGs from the cervical to the lumbar level were carefully dissected on ice under a microscope in Ca^2+/^Mg^2+^ free Hank’s balanced salt solution (HBSS). Media was replaced with collagenase II (200 U/mL, Worthington, LS004177)/dispase II (3.75 U/mL, Sigma-Aldrich, D4693) solution and incubated in 37 °C for 40 min, then underwent shaking (70 r.p.m, Fisherbrand™ 3D Rotators, 88-861-047) for 15 min at room temperature. DRGs were dissociated using prepared sterile flame polished pipettes, and filtered at 100 μm (BD Falcon, 352360). Cells were collected at 380 g and washed with ice-cold CMF-HBSS twice. DRGs cultures were grown at 37 °C in Neurobasal-A medium supplemented with 2% B-27 and 1% Glutamax-I (Gibco).

#### Primary rat hippocampal neurons culture

Primary neurons were isolated from the hippocampus of postnatal day 1 Sprague-Dawley rats and prepared according to an established protocol with minor modifications^94^. Briefly, after the removal of the remaining meninges and midbrain tissue, hippocampi were excised from the cortex in ice-cold HBSS and then subjected to 0.1% Trypsin at 37 °C for 15 min. The digestion reaction was terminated by DMEM media containing 10% FBS, then the digested tissue was dispersed with DNase I (20 U/mL, Thermo Scientific, EN0521), triturated with fire-polished Pasteur pipettes, and filtered through a 70 μm strainer (BD Falcon, 352350). Cells were plated with Neurobasal-A medium containing 2% B-27 and 1% Glutamax-I. Half of the medium was refreshed every two days.

#### Primary rat astrocytes culture

Astrocytes were harvested as previously described^12^. P3 Sprague Dawley rats were decapitated, and the cortices were removed, finely minced, and treated with 0.1% trypsin in EDTA at 37 °C for 15 min. After this digestion process, the tissue was washed with DMEM media (containing 10% FBS and 20 U/mL DNase I) and mechanically dissociated with fire-polished Pasteur pipettes. The cell suspension was filtered through a 70 μm strainer and centrifuged at room temperature at 300 g for 8 min. The resultant pellet was resuspended in DMEM with 10% FBS. The primary astrocytes used for all experiments were between passages 3 and 5.

#### Primary rat meningeal fibroblasts culture

Meningeal fibroblasts were simultaneously prepared during the dissociation of DRGs and harvested as previously described^46^. Meninges were carefully removed from the surface of the spinal cord using forceps, minced into small pieces, and subjected to enzymatic digestion with 200 U/mL of collagenase II and 3.75 U/mL of dispase II for 40 min at 37 °C. Meninges were triturated and cells were filtered through a 70 μm strainer, washed, and centrifuged (360 g for 8 min) to obtain single-cell suspensions. Meningeal fibroblasts were maintained in DMEM with 10% FBS and were used for all experiments between passages 3 and 5.

#### Collagen gradient crossing assay

Collagen gradients were prepared similar to those used to study the inhibitory properties of CSPGs as previously described^12,34^. Pre-sterilized glass coverslips (12 mm diameter, Chemglass, CLS-1760-012) were rinsed with sterile water and then coated with poly-L-lysine hydrobromide (0.1 mg/mL, Sigma-Aldrich, P6282) overnight at room temperature or at 37 °C for 3 h. After allowing the coverslips to air dry for 1 hour and rinsing them with sterile water three times, prepared nitrocellulose was applied onto the PLL side and left to dry for 10 min. Collagens I (Corning, 354236), III ( Advanced BioMatrix, 5021), IV (Advanced BioMatrix, 5022), VI (Rockland, 009-001-108), Aggrecan (Sigma-Aldrich, A1960), or fibronectin (Corning, 354008) with natural laminin (5 μg/mL, Gibco, 23017-015 ) in HBSS were dried onto the coverslip surface to create the spot gradients. The spots were then bathed with laminin (5 μg/mL) in HBSS at 37 °C for at least 2 h. Dissociated DRGs from one adult female rat were aliquoted among 14 wells of a 24-well plate and incubated for 5 days at 37 °C. Dissociated hippocampal neurons were plated at a density of 10,000/cm^2^ and incubated for 7 days. Primary astrocytes and meningeal fibroblasts were plated at a density of 2,000/cm^2^ and incubated for 3 days. For the drug screening experiments, appropriate concentrations of the following compounds were utilized: ISP^12^, Ixabepilone^46^ (MedChemExpress, HY-10222), Hamster Anti-Rat Itgb1^40^ (BD Pharmingen, 555003), Nilotinib (MedChemExpress, HY-10159), Sitravatinib (MedChemExpress, HY-16961), Merestinib (MedChemExpress, HY-15514), recombinant ΔPLL domain (A26-N175)^95^ of rat GPR56 (rΔPLL-rbFc, this paper), Dihydromunduletone (MedChemExpress, HY-101483), Y-27632 dihydrochloride (LC laboratories, Y-5301), Ruthenium red (MedChemExpress, HY-103311), EGTA (Sigma-Aldrich, E8145), and GsMTx4 peptide (MedChemExpress, HY-P1410). These compounds were incubated with DRGs for 1 h before plating and were refreshed every 24 h. For enzyme degradation assay, different concentrations of collagenase II (Worthington) from 0.01-100 U/mL were added to coverslips for 2 h after laminin bath. The coverslips were washed three times with HBSS before cells plating.

#### Immunofluorescence

Coverslips were fixed in 4% paraformaldehyde for 10 min, washed with PBS twice, permeabilized and blocked in EveryBlot Blocking Buffer (Bio-Rad, 12010020) containing 0.3% Triton X-100, then stained in the same blocking buffer. The following primary antibodies were used for staining: mouse anti-collagen I (1:500, Invitrogen, MA1-26771), rabbit anti-collagen I (1:60, Bio-Rad, 2150-1908), mouse anti-collagen III (1:100, Santa Cruz, sc-271249), rabbit anti-collagen IV (1:500, Abcam, ab6586), mouse anti-collagen IV (1:50, Developmental Studies Hybridoma Bank, M3F7), goat anti-collagen VI (1:100, SouthernBiotech, 1360-08), rabbit anti-Laminin (1:500, Sigma-Aldrich, L9393), mouse anti-βIII Tubulin (1: 1,000, Promega, G712A), chicken anti-βIII Tubulin (1:1,000, Aveslabs, TUJ), mouse anti-CSPG (1:200, Sigma-Aldrich, C8035), mouse anti-fibronectin (1:200, BD Biosciences, 610077), rabbit anti-β-Amyloid Precursor Protein (CT695) (1:500, Invitrogen, 51-2700), mouse anti-β-Amyloid (6E10) (1:100, Biolegend, 803004), mouse anti-synaptic vesicle glycoprotein 2 (SP2/0) (1:50, Developmental Studies Hybridoma Bank, SV2), mouse anti-Tau-1 (PC1C6) (1:100, Sigma-Aldrich, MAB3420), rabbit anti-V-Glut1 (1:100, Sigma-Aldrich, V0389), rabbit anti-Neurofilament 200 (1:200, Sigma-Aldrich, N4142), chicken anti-GFAP (1:400, Millipore, AB5541), mouse anti-α-tubulin (1:100, Developmental Studies Hybridoma Bank, 12G10).

Coverslips were rinsed in TBS 3 times and incubated with secondary antibodies (Jackson ImmunoResearch) diluted in blocking buffer at 1:400 for 1 h at room temperature. Nuclei were stained with DAPI (1:1,000, Sigma-Aldrich, D9542) and coverslips were mounted using Antifade Mounting Media (4 μL per coverslip, Vectashield, H-1000). Images were captured using an Axio Imager M2 microscope (Zeiss, Germany) equipped with an ORCA-Flash4.0 V3 Digital CMOS camera (Hamamatsu, C13440-20CU, Japan), or an 800 confocal laser-scanning microscope (Zeiss, Germany).

#### Expression of recombinant rΔPLL-rbFc protein

In the extracellular region of GPR56, a Pentraxin/Laminin/neurexin/sex-hormone-binding-globulin-Like (PLL) domain was identified, which directly engages the GPR56 ligand to initiate G protein signaling^95^. Rabbit IgG1 Fc2-tagged ΔPLL (A26-N175 from rat GPR56) fusion proteins, termed rΔPLL-rbFc, were designed and used for blocking assay. General procedures of plasmid construction and protein purification has been previously described in detail. Briefly, recombinant rΔPLL-rbFc was expressed in HEK-293t cells by transiently transfection. The culture media was changed to serum-reduced Opti-MEM (Gibco, 31985070) 24 h after transfection. Conditioned media was collected 72 h later and concentrated (Millipore, 10000MW). rΔPLL-rbFc was purified through protein A-agarose (Santa Cruz Biotechnology, sc-2001).

#### Immunoblotting

Meningeal fibroblasts were passaged and placed in culture for 12 h. Then, different concentrations of aspirin, pirfenidone, and an equal volume of vehicle were added. After 48 h, cells were washed with ice-cold PBS twice and lysed with RIPA lysis buffer (Thermo, 89900) containing a protease inhibitor cocktail (Thermo, 78430) for 10 min in 4 °C. Cells lysates were scraped and clarified by centrifugation at 13,000 g for 10 min at 4 °C. Lysates supernatant with 6 × Laemmli SDS buffer (Thermo, J61337-AD) was boiled for 10 min at 97°C and loaded into 8% SDS-PAGE gels with Precision Plus Protein Standard (Bio-Rad, 161-0374/161-0375). Separated proteins were transferred to a nitrocellulose membrane (0.2 μm, Bio-Rad, 1620112) at 85 V for 120 min under an ice-water bath. The membrane was blocked at room temperature with EveryBlot Blocking Buffer (Bio-Rad, 12010020) for 30 min before incubating with an antibody against Collagen I (1:500, Bio-Rad, 2150-1908), Collagen III (1:300, Santa Cruz, sc-271249), or β-Actin (1:1,000, Cell Signaling, 3700) at 4 °C overnight. After three 5-min washes with TBS/T, membranes were incubated with a 1:2,000 dilution of anti-mouse/rabbit IgG, HRP-linked secondary antibodies (Cell Signaling, 7076/7074) at room temperature for 1.5 h. After washing three times with TBS/T and ddH_2_O, the blot image was developed using a Clarity Western ECL Substrate (Bio-Rad, 170-5060) and scanned by Odyssey Fc Imager (LI-COR, OFC-0741). Band intensities shown in immunoblots were quantified using Image J software (NIH, 1.52p version) and normalized to the protein levels of the β-Actin control. All experiments were repeated three times independently.

### In vivo assay

#### Animals

Female C57BL6/J mice and Sprague Dawley (SD) rats were obtained from Inotiv or Charles River. Animals were housed in an environment of 22 °C and 12 h light/dark cycle with free access to food and water ad libitum. All experiments were approved by the Institutional Animal Care and Use Committee of Case Western Reserve University (CWRU), Cleveland. For thoracic spinal cord injury, 20 g body weight mice and 230-250 g body weight rats were used. For dissection and culture of rat dorsal root ganglion neurons, 170-190 g body weight rats were used.

#### Surgery

Spinal cord crushes and contusions were performed as previously described^1,12,27,46^. In brief, animals were anaesthetized with isoflurane and T10 was exposed by laminectomy. 2s crush injuries of different severities (incomplete and complete) were performed by inserting spacers into Dumont forceps No. 5 (Fine Science Tools, 11251-20), resulting in tip widths of 0.5 mm or 0.2 mm and spacing of 0.25 mm (incomplete injury model) or 0 mm (without spacers, complete injury model) upon closure. Rat with cord injuries were anesthetized by intraperitoneal injection of ketamine (75-100 mg/kg) and xylazine (5-10 mg/kg). The vertebral column was secured by clamping the T9 and T11 vertebral bodies using forceps attached to the base of an Infinite Horizon Impact Device (IH-0400). A 2.5 mm stainless steel impactor tip was positioned over the midpoint of T10 and impacted with forces of 250 kdyn force. Any animals displaying abnormal displacement/force graphs or deviating more than 10% from the setting force (Impactor Application v5.0.4) were promptly excluded from the study. After surgery, muscles and fascia layers were sutured with 4-0 (mice) or 3-0 (rats) absorbable sutures, and the skin was closed with wound clips. Animals were allowed to recovered on a warming pad and given meloxicam (10 mg/kg *s.c.*, every 12 h up to 72 h) for pain which we monitored closely. Manual bladder emptying was conducted 2-3 times daily. Animals were randomly grouped and assigned numbers, and were treated blind to experimental conditions.

#### Drug preparation and treatment

The amount of drugs for one weeks worth of injections was prepared and packaged in DMSO and stored under -20 °C conditions, with each use at 37 °C for a complete dissolution. The diluent for pirfenidone includes 30% PEG300, 5% Tween-80, and 60% sterile saline, and the concentration of DMSO is maintained at 5%. The diluent for aspirin includes 15% PEG300, 2.5% Tween-80, and 80% sterile saline, and the concentration of DMSO is maintained at 2.5%. For treatment, rats were injected with aspirin (500 μL *i.p.*, 45 mg/kg, one time a day, 8 dpi onward) and pirfenidone (500 μL *i.p.*, 16 mg/kg, three times a day, 1-7 dpi; 32 mg/kg, three times a day, 8-14 dpi; 48 mg/kg, three times a day, 15 dpi onward) or vehicle control. Metergoline (500 μL *i.p.*, 7.5 mg/kg) was administered to uninjured rats and combination-treated responders.

#### Lesion analysis and quantification

Sections stained for collagen I, collagen III, PDGFRβ, CSPG, laminin, CD68, and GFAP were scanned at constant exposure settings. Images were converted and thresholded (Intermodes, Dark background) in Fiji (NIH, ImageJ 1.52p). Immunostaining intensity were measured and normalized to the intact area.

#### RNA-seq and bioinformatic analysis

After a quick ice-cold PBS perfusion, spinal cords were dissected quickly and 0.5 mm long pieces of tissue centered on the injury core were carefully dissociated in ice-cold PBS. Tissues were directly lysed in ice-cold RNA isolater (Vazyme, R401) for total RNA isolation using RNeasy Mini Kit (Qiagen, 74004). Each group contains three replicates. RNA-seq for mice tissues were carried out by GENEWIZ (Project No. 80-767360528). In brief, total RNA of each sample was quantified and qualified by Agilent 2200 Bioanalyzer (Agilent Technologies). 1 μg total RNA was used for the ensuing library preparation. Next generation sequencing library preparations were constructed according to the manufacturer’s protocol. Sequencing was carried out using a 2 x 150 paired-end (PE) configuration. We processed the pass filter data in FASTQ format using Cutadapt (V1.9.1) to make it high-quality clean data. Reads were aligned via software Hisat2 and an average uniquely aligned rate of 87.74% was obtained. Read counts were generated by HTSeq v0.6.1. Differential expression analysis was performed via a DESeq2 Bioconductor package. RNA-seqs for rat tissues were carried out by BGI Genomics (Project No. F23A480000612_RATgsvoR) using DNBSEQ platform, and the average mapping ratio of genes was 74.59%. Genes with an adjusted *P* value less than 0.05 and log-transformed fold change larger than 0.25 were selected for bioinformatic analysis (575 downregulated gene and 331 upregulated gene, aspirin and pirfenidone treated group vs vehicle treated group). GO enrichment analyses were performed using DAVID (NIH, v2023q2).

#### Tracing

To label rats CST axons, AAV8-hSyn-EGFP (Addgene, 50465) was injected into the sensorimotor cortex at 14 dpi. Virus was injected at 8 sites (1.0 μL each at a rate of 0.2 μL/min): ± 1.6 mm and ± 2.6 mm lateral; 1.3 mm and 2.3 mm posterior to bregma; 1.5 mm deep. To label rats propriospinal neurons, 10% biotinylated dextran amine (BDA) 10,000 (Invitrogen, D1956) was injected into the gray matter of spinal cords at 28 dpi (one and two segments rostral to SCI lesions). BDA was injected at 4 sites (0.5 μL each at a rate of 0.1 μL/min): ± 0.4 mm lateral; 2 mm and 3 mm rostral to SCI lesions; 1.1 mm deep.

## Spinal cord stereotaxic injection of collagen

Rats were anaesthetized with ketamine (75-100 mg/kg) and xylazine (5-10 mg/kg), and surgeries were performed under aseptic conditions. T9 and T11 of the spine were fixed with forceps and suspended from the fixation plate. Collagens (0.7 mg/mL) were injected into two sites (0.5 mm bilateral injection, 1.2 μL/site) 1.8 mm below the surface at the T10 segment spinal cord segment (0.2 μL/min) using glass micropipettes connected to an injector (Nanoliter) under the control of a microprocessor-based controller (SYS-Micro4). The needle was left in place for 5 min before removing. Rats received meloxicam (10 mg/kg *s.c.*, every 12 h up to 72 h) for pain relief, and were perfused at 7 days after injection. 30 µm thick horizontal sections of the spinal cord were collected for immunohistochemistry analysis.

### Perfusion, sectioning, and immunohistochemistry

Mice or rats were given a lethal dose of anesthesia and transcardially perfused with 4% paraformaldehyde in PBS for 10 min. Spinal cords were dissected out, post-fixed in 4% paraformaldehyde overnight at 4 °C, and cryoprotected with 30% sucrose in PBS for 3 days at 4 °C. After embedding into O.C.T mounting media and frozen in dry ice, serial sections were collected on a LEICA cryostat (CM-3050-S) at 20 μm (mice) or 30 μm (rats) thicknesses, dried on a 37 °C warming plate, and stored at -80 °C until processed. Before staining, sections were warmed to room temperature and treated with EveryBlot Blocking Buffer (Bio-Rad, 12010020) containing 0.5% Triton X-100 for 1 h at room temperature. After blocking, slides were subjected to primary antibodies diluted in EveryBlot Blocking Buffer overnight at 4 °C. The following primary antibodies were used with the indicated dilutions: goat anti 5-HT (Serotonin) (1:5,000, Immunostar, 20079), rabbit anti-collagen I (1:60, Bio-Rad, 2150-1908), rabbit anti-collagen I (1:200, Cell Signaling, 72026), rabbit anti-collagen III (1:60, Bio-Rad, 2150-1948), mouse anti-collagen III (1:100, Santa Cruz, sc-271249), rabbit anti-collagen III (1:100, LS Bio, LSLJC352027), rabbit anti-collagen IV (1:500, Abcam, ab6586), rabbit anti-collagen VI (1:200, Abcam, ab182744), chicken anti-GFAP (1:400, Millipore, AB5541), mouse anti-CSPG (1:200, Sigma-Aldrich, C8035), goat anti-Iba1 (1:100, Abcam, ab5076), rat anti-PDGFRb (1:100, Invitrogen, 14-1402-82), rabbit anti-Laminin (1:500, Sigma-Aldrich, L9393), mouse anti-CD68 (1:100, Bio-Rad, MCA341), rabbit anti-beta-amyloid precursor protein (CT695) (1:250, Invitrogen, 51-2700), Rabbit anti-NG2 chondroitin sulfate proteoglycan (1:100, Sigma-Aldrich, AB5320). The next day, sections were washed with TBS and incubated in the appropriate Alexa Fluor 488-, 594- or 647-conjugated secondary antibody (Jackson ImmunoResearch) at room temperature for 2 h. DAPI (1:1,000 in PBS) was used for nucleic acid staining. All sections of spinal cord were screened with a Axio Imager M2 microscope (Zeiss, Germay) or a confocal laser-scanning microscope (Zeiss 800, Germay).

### 5-HT density quantification

Sagittal spinal cord sections stained with 5-HT antibody (1:5000, Immunostar, 20079) were scanned under constant exposure settings. All Images were converted and thresholded (Intermodes, Dark background) in Fiji (NIH, ImageJ 1.52p). A series of 1 x 1 mm^2^ rectangles were superimposed on the gray matter in sequence, from the distal border to the lumbar spinal cord. Densities were computed within each rectangle as the ratio of 5-HT fibers (normalized to 1 x 1 mm^2^ rectangle on proximal border). The average ratio of 4-6 images per rat were used for quantification.

### Behavioral analysis

In contusive SCI and complete SCI experiments on rats, the Basso, Beattie and Bresnahan (BBB) locomotor scale was conducted by blinded observer on day 1, 7 and weekly thereafter through week 9 as previously described. The Basso Mouse Score (BMS) was performed on mice in ISP treatment experiments. To evaluate differences in incomplete and complete SCI experiments, a horizontal ladder-walking test, with clear plexiglass side walls (1.1 m in length) and evenly spaced (2 cm) sterile wooden rods elevated 18 cm from the ground, was executed to capture hindlimb performance on days 0 and, 14 and weekly thereafter until week 9. To prevent muscle memory, rats did not undergo pre-training. A complete miss, slip, or replacement of the paw during placement was considered as an error. During ladder-walking, the gait was captured by a camera fixed at a side position. We installed a custom-made floor onto a horizontal ladder to facilitate the camera recording of hindlimb gaits of rats during overground walking.

### Statistical analysis

All statistical analyses were performed in GraphPad Prism (USA). All data underwent normality testing (D’Agostino & Pearson test and Anderson-Darling test) in advance. Two-tailed unpaired Student’s *t*-test post-hoc Welch’s correction and one-way ANOVA post-hoc Bonferroni correction was used when data were distributed normally. Otherwise, Mann-Whitney test or Kruskal-Wallis test post-hoc Dunn correction were performed. Two-way ANOVA post-hoc Bonferroni or Dunnett correction were applied for multiple comparison procedures. The significance level was set as *P <* 0.05. All statistical results are provided as Source Data.

## Acknowledgements

We thank Jingqiang (Bruce) You for animal care and independent behavioral analysis. Financial support was provided by The Brumagin-Nelson Fund, The Griffin Family, The Timothy Brodigan Trust, The Kaneko Family Fund and NIH NINDS grant (1R01NS122371). We gratefully acknowledge the major support of the Government of Canada’s New Frontiers in Research Fund (Mend the Gap, No. NFRFT-2020-00238), and the National Natural Science Foundation of China (No. 82201533).

## Author contributions

Y.B. and J.S. conceived the project. Y.B. and W.D. conducted experiments and analyzed the data. Y.B., W.D., and J.S. wrote the manuscript.

## Data availability

The data are available from the corresponding author upon reasonable request.

