## Supporting Information for "Collagen I is a critical organizer of scarring and CNS regeneration failure"

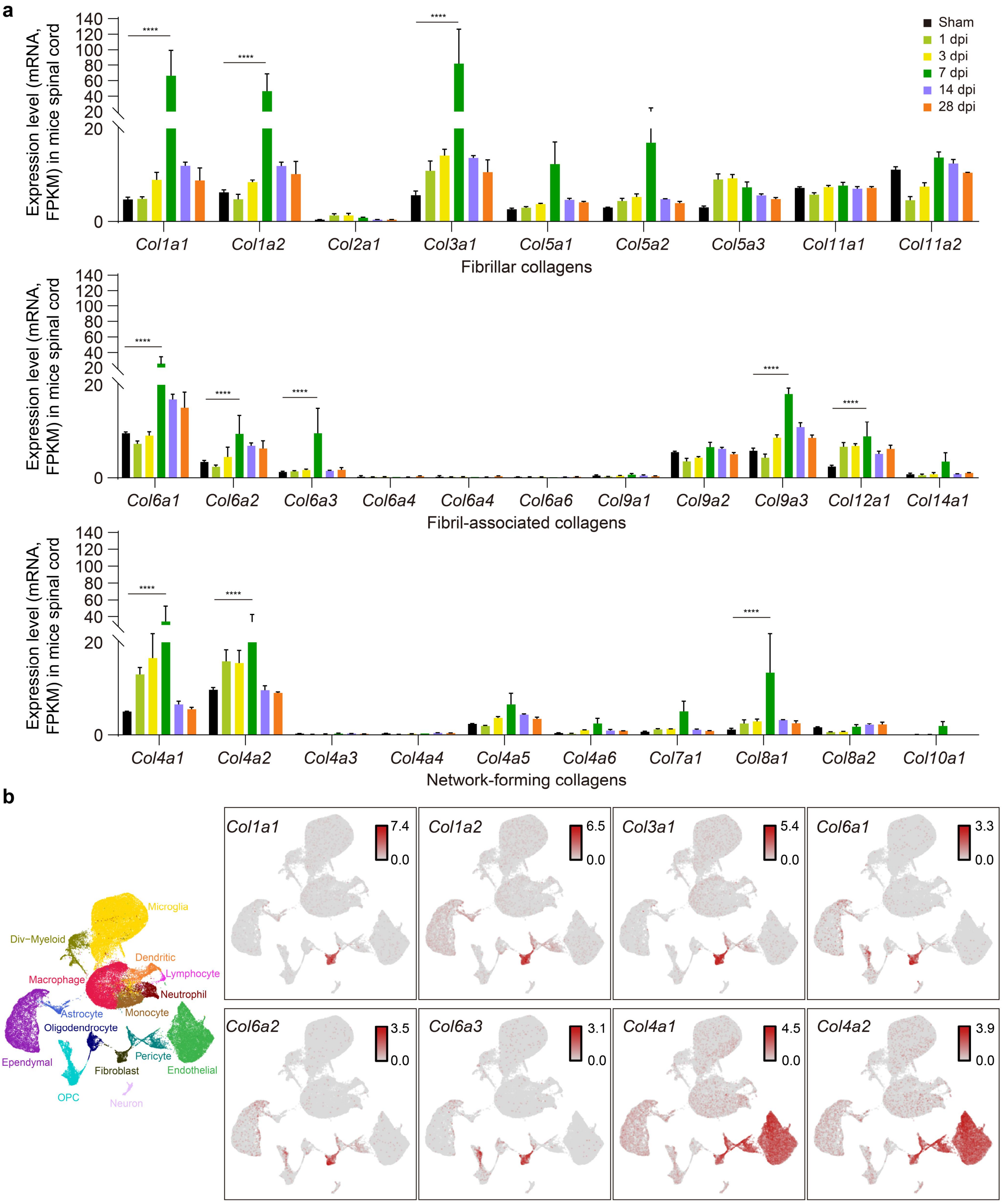

**Extended Data Fig. 1 | Characterizing the expression profiles of various collagen types in the spinal cord following injury**.
**a**, Expression of genes associated with fibrillar collagens, fibril-associated collagens, and network-forming collagens in adult mice in sham animals, and at 1, 3, 7, 14, and 28 dpi, using bulk RNA-seq. FPKM, fragments per kilobase of transcript sequence per million mapped fragments. Data shown as mean ± s.e.m ($n$ = 3 per group). ****$p < 0.0001$, Two-way RM ANOVA followed by post-hoc Dunnett correction.
**b**, Gene expression patterns of *col1a1, col1a2, col3a1, col4a1, col4a2, col6a1, col6a2,* and *col6a3* in different cell clusters identified in UMAP plots. *Col I* (*col1a1, col1a2*), *Col III* (*col3a1*) and *Col VI* (*col6a1, col6a2, col6a3*) expressions are highly enriched in fibroblasts. *Col IV* (*col4a1, col4a2*) expression is highly enriched in fibroblasts, pericytes, and endothelial cells. UMAP plot representing all cells collected from both the uninjured spinal cord and the injured spinal cord at 1, 3, and 7 dpi, totaling 66,178 cells. Values indicate log-normalized counts per cell. Raw data from NCBI GEO repository (no. GSE162610), and visual online tools (*https://jaeleelab.shinyapps.io/sci_singlecell/*)[1].

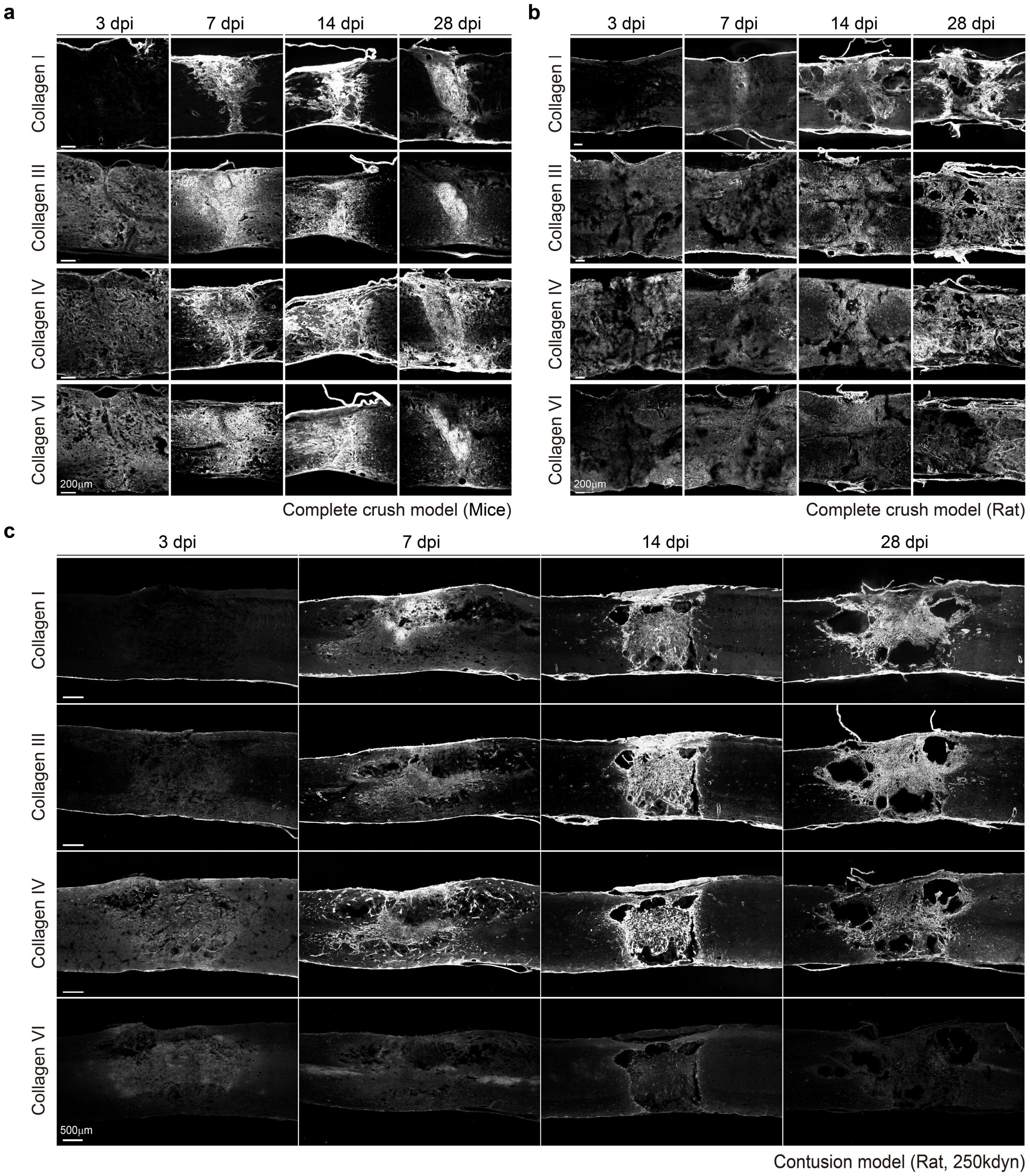


**Extended Data Fig. 2 | Characterizing the expression profiles of various collagen types in the spinal cord following injury**. **a**, **b**, **c** Images of spinal cord sections stained with antibodies against collagen I, III, IV and VI at different time points after injury in adult mice and rats. Scale bars, 200 μm (**a**, **b**), 500 μm (**c**).

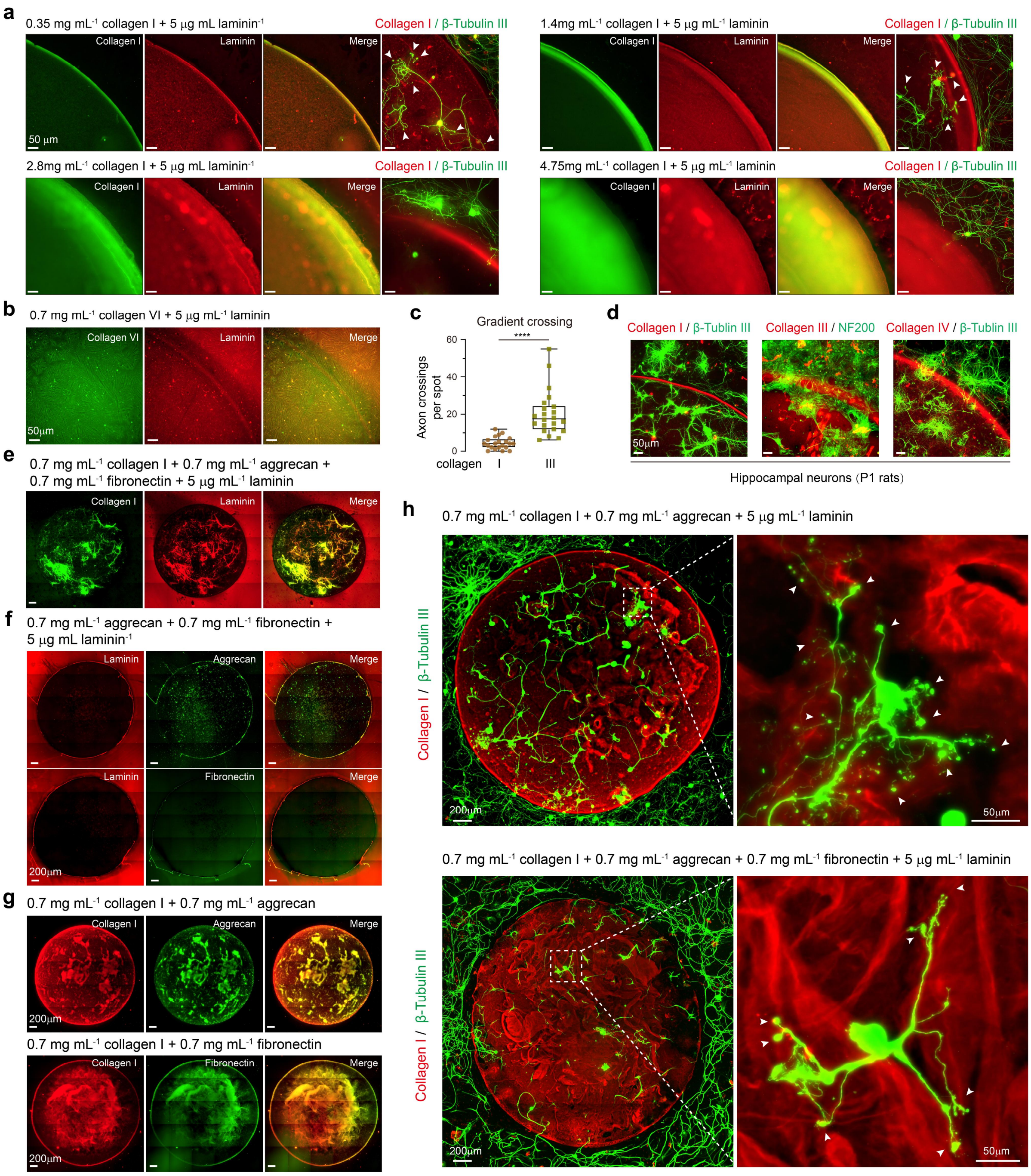
a
0.35 mg mL-1 collagen I + 5 µg mL laminin-1
Collagen I / β-Tubulin III
Collagen I
Laminin
Merge
50 µm
1.4mg mL-1 collagen I + 5 µg mL-1 laminin
2.8mg mL-1 collagen I + 5 µg mL laminin-1
4.75mg mL-1 collagen I + 5 µg mL-1 laminin
b
0.7 mg mL-1 collagen VI + 5 µg mL-1 laminin
Collagen VI
50µm
c
Gradient crossing
Axon crossings per spot
****
collagen
I
III
d
Collagen I / β-Tublin III
Collagen III / NF200
Collagen IV / β-Tublin III
Hippocampal neurons (P1 rats)
e
0.7 mg mL-1 collagen I + 0.7 mg mL-1 aggrecan + 0.7 mg mL-1 fibronectin + 5 µg mL-1 laminin
f
0.7 mg mL-1 aggrecan + 0.7 mg mL-1 fibronectin + 5 µg mL laminin-1
Aggrecan
Fibronectin
200µm
g
0.7 mg mL-1 collagen I + 0.7 mg mL-1 aggrecan
0.7 mg mL-1 collagen I + 0.7 mg mL-1 fibronectin
h
0.7 mg mL-1 collagen I + 0.7 mg mL-1 aggrecan + 5 µg mL-1 laminin
Collagen I / β-Tubulin III
0.7 mg mL-1 collagen I + 0.7 mg mL-1 aggrecan + 0.7 mg mL-1 fibronectin + 5 µg mL-1 laminin

**Extended Data Fig. 3** | ***In vitro* simulation of fibrous scar ECM formation**.

**a**, Increasing collagen I concentration modifies the collagen I gradient and inhibits DRG growth either in the center or across the rim.

**b**, Collagen VI-laminin mixture failed to form gradient spots. Two commercial products were tested (Rockland, No.: 009-001-108; SouthernBiotech, No.: 1300-02).

**c**, Quantification of the number of axons crossing the rims of collagen I (0.7 mg/mL)-laminin (5 mg/mL) and collagen III (0.7 mg/mL)-laminin (5 mg/mL) spots, related to Fig. 1f ($n \geq 21$). ****$p < 0.0001$, Mann Whitney test. Data are min to max, show all points.

**d**, Immunostained images of hippocampal neurons (P1 rats) across the collagen I/III/IV spot rim. Scale bars, 50 μm.

**e**, Images of the collagen I-aggrecan-fibronectin-laminin mixture spot showing the distribution of collagen I and laminin. Four mixtures form a fiber structure.

**f**, Images of the aggrecan-fibronectin-laminin mixture spot showing the distribution of each component. Three mixtures failed to form the fiber structure.

**g**, Images of the collagen I-aggrecan and collagen I-fibronectin mixture showing the distribution of each component. Both mixtures can form the fiber structure.

**h**, Immunostaining of β-tubulin III (green) in adult DRGs on the collagen I-aggrecan-laminin mixture and collagen I-aggrecan-fibronectin-laminin mixture spot. The white arrows indicate the appearance of endballs on neurites. Scale bars, 200 μm (**e**, **f**, **g**, **h**).

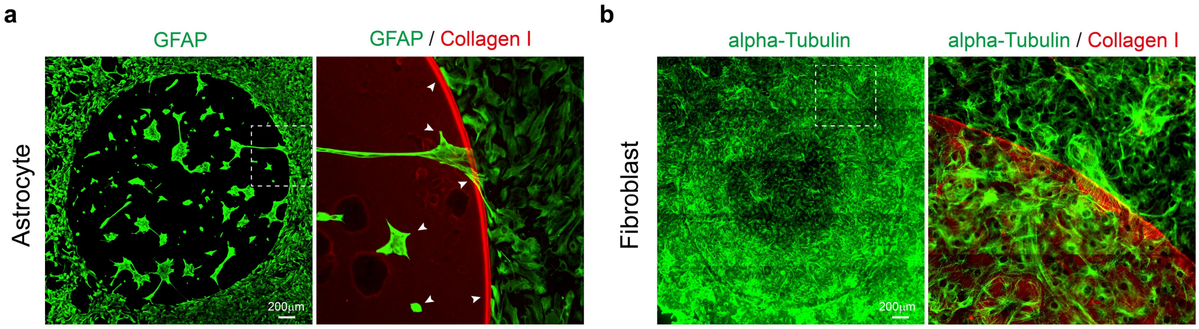


**Extended Data Fig. 4 | Astrocyte and fibroblast responses to collagen I gradient spot**.
**a**, Immunostaining showing the behavior of purified GFAP-positive astrocytes (green) on collagen I gradient spot (red).
**b**, Immunostaining showing behavior of purified meningeal fibroblasts on collagen I gradient spot (red). Display of fibroblast cytoskeleton using anti-alpha-Tubulin, green. Scale bars, 200 μm (**a**, **b**).

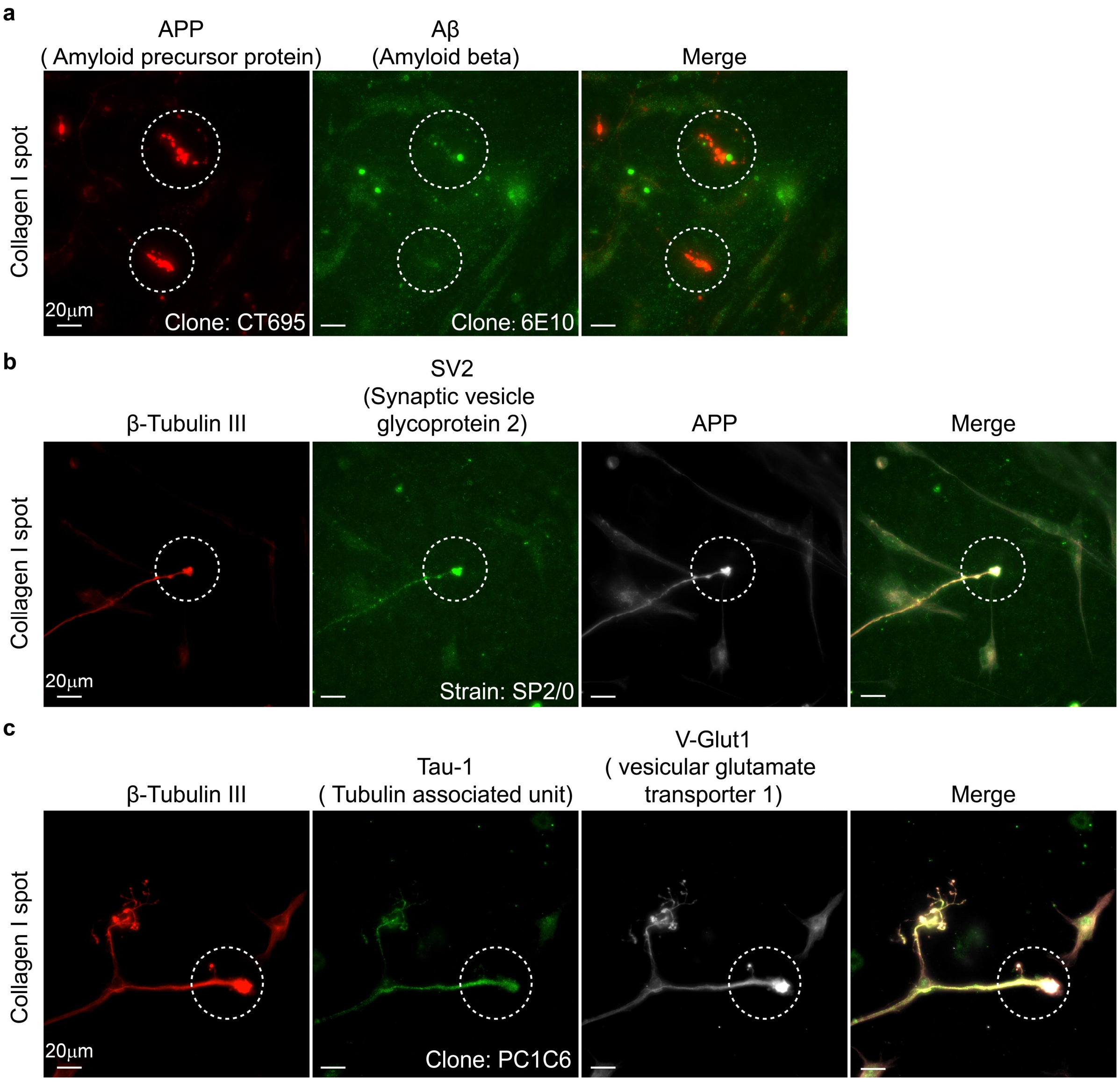


**Extended Data Fig. 5 | The characteristics of "dystrophic endballs" in DRG axons on collagen I gradient**.

**a**, Immunostaining indicates that APP (red) is present in these "dystrophic endballs" and has not undergone further hydrolysis into Aβ (green).

**b**, Immunostaining of the presynaptic membrane marker SV2 in "dystrophic endballs" shows positive signals, implying that these endballs might resemble a synaptic-like structure.

**c**, Representative image of "dystrophic endballs" immunolabelled for β-tubulin III (red), Tau-1 (green), and V-Glut1 (white). Scale bars, 20 μm (**a**, **b**, **c**).

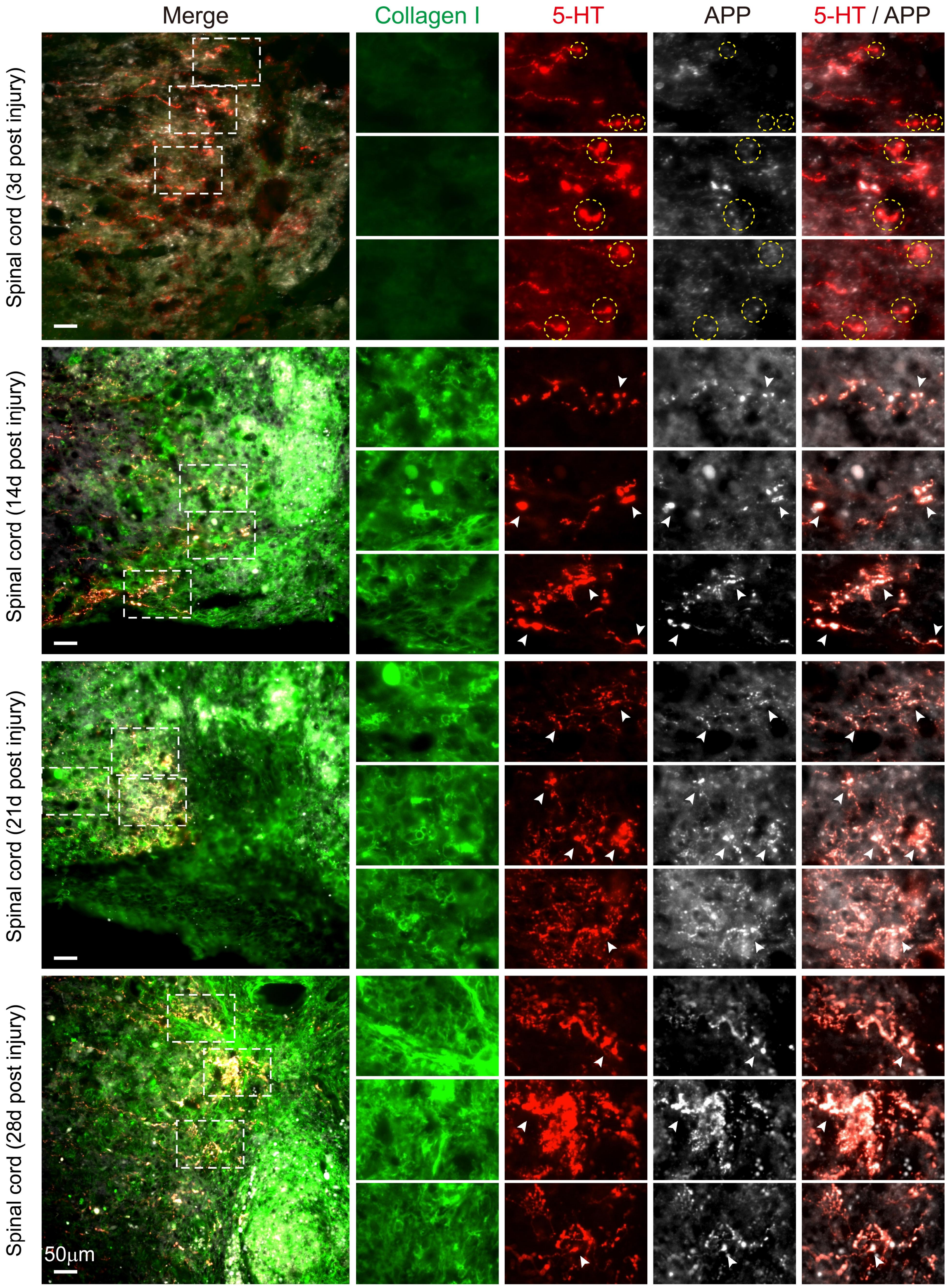

Merge
Collagen I
5-HT
APP
5-HT / APP
Spinal cord (3d post injury)
Spinal cord (14d post injury)
Spinal cord (21d post injury)
Spinal cord (28d post injury)
50μm

**Extended Data Fig. 6 | APP accumulates in the endballs of serotonergic axons in the vicinity of collagen I after SCI**. Representative microscopic images of collagen I (green), 5-HT (red), and APP (white) expression around the lesion sites at different time points in SCI mice. 3 days after SCI, collagen I has not yet been detected, and at this point, few $APP^+$ endballs could be observed (yellow circles indicated representative regions). From 7-28 days, collagen I begins to accumulate significantly in the lesion site, and a substantial aggregation of APP was observed in the endballs (white arrows). Scale bars, 50 μm.

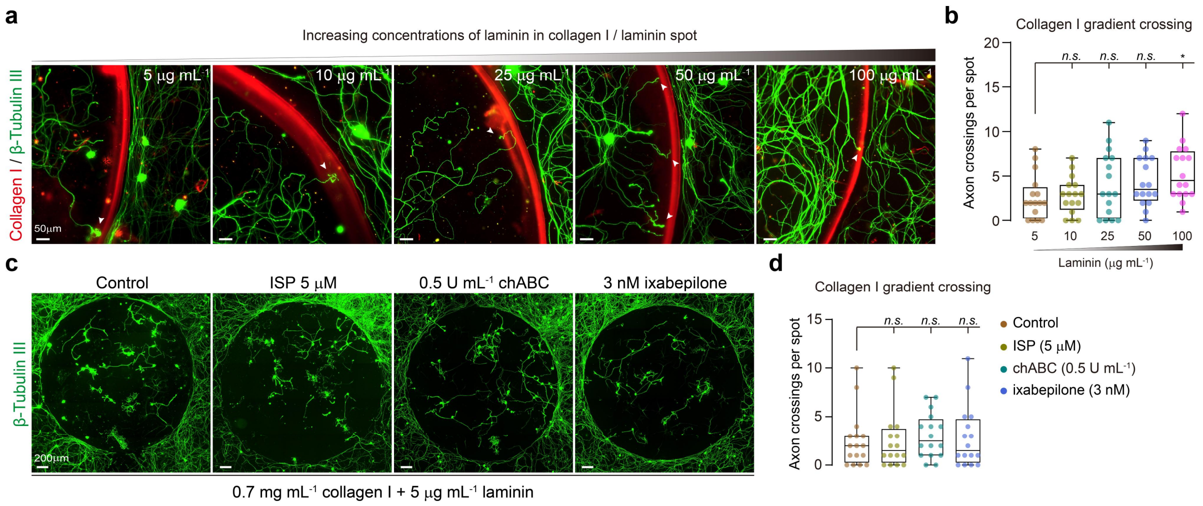


**Extended Data Fig. 7** | **Screening strategies to overcome retraction induced by collagen I**.

**a**, The impact of laminin concentration gradients on neurite outgrowth within collagen I spots.

**b,** Quantification of numbers for DRG axons crossing the inhibitory rim according to **a** ($n$ = 16 per group). *$p$ = 0.0209, *n.s.*, not significant. Kruskal-Wallis test followed by post-hoc Dunn correction.

**c,** Representative microscopy images of β-tubulin III (green) in DRGs on a complete collagen I gradient spot with added ISP, chABC, and ixabepilone.

**d,** Quantification of numbers for DRG axons crossing the inhibitory rim according to **c** ($n$ = 16 per group). *n.s.*, not significant. Kruskal-Wallis test followed by post-hoc Dunn correction. Data are min to max, show all points. Scale bars, 50 μm (**a**), 200 μm (**c**).

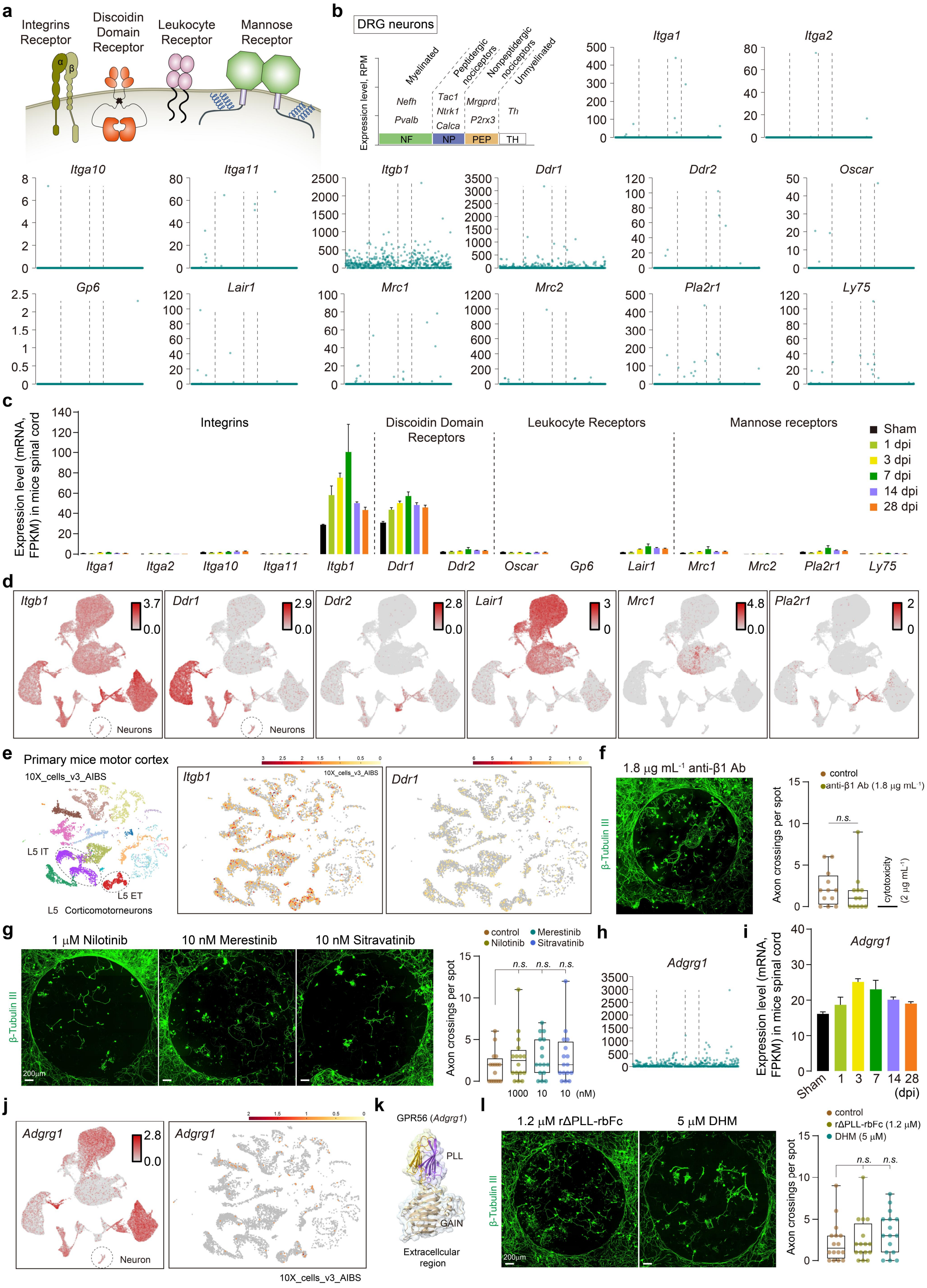

a
Integrins Receptor
Discoidin Domain Receptor
Leukocyte Receptor
Mannose Receptor
α
β
b
DRG neurons
Expression level, RPM
Myelinated
Peptidergic nociceptors
Nonpeptidergic nociceptors
Unmyelinated
Nefh
Pvalb
Tac1
Ntrk1
Calca
Mrgprd
P2rx3
Th
NF
NP
PEP
TH
Itga1
Itga2
Itga10
Itga11
Itgb1
Ddr1
Ddr2
Oscar
Gp6
Lair1
Mrc1
Mrc2
Pla2r1
Ly75
c
Expression level (mRNA, FPKM) in mice spinal cord
Integrins
Discoidin Domain Receptors
Leukocyte Receptors
Mannose receptors
Sham
1 dpi
3 dpi
7 dpi
14 dpi
28 dpi
d
Neurons
e
Primary mice motor cortex
10X_cells_v3_AIBS
L5 IT
L5 ET
L5 Corticomotorneurons
f
1.8 μg mL⁻¹ anti-β1 Ab
β-Tubulin III
control
anti-β1 Ab (1.8 μg mL⁻¹)
Axon crossings per spot
n.s.
cytotoxicity (2 μg mL⁻¹)
g
1 μM Nilotinib
10 nM Merestinib
10 nM Sitravatinib
200μm
control
Nilotinib
Merestinib
Sitravatinib
1000
10
10 (nM)
h
Adgrg1
i
Sham
1
3
7
14
28
(dpi)
j
Neuron
k
GPR56 (Adgrg1)
PLL
GAIN
Extracellcular region
l
1.2 μM rΔPLL-rbFc
5 μM DHM
rΔPLL-rbFc (1.2 μM)
DHM (5 μM)

**Extended Data Fig. 8 | Expression of collagen receptors**.

**a**, Schematic diagram of collagen receptors.

**b**, Single-cell absolute expression levels (reads per million, RPM) of collagen receptors in dissociated DRGs (Schematic diagram adapted from Dmitry Usoskin, *et al*. 2014)[2]. Raw data from NCBI GEO repository (no. GSE59739), and visual online tools (http://linnarssonlab.org/drg/).

**c**, Expression of collagen receptors in adult mice in sham animals, and at 1, 3, 7, 14, and 28 dpi, using bulk RNA-seq. FPKM, fragments per kilobase of transcript sequence per million mapped fragments. Data shown as mean ± s.e.m ($n$ = 3 per group).

**d**, UMAP pattern of the collagen receptors in spinal cord. Annotated cell type described in Extended Data Fig. **1b**[1].

**e**, Expression pattern of Integrin-β1 (*Itgb1*) and DDR1 (*Ddr1*) in adult mouse primary motor cortex neurons. UMAP projection based on analysis of 10x_cells_v3_AIBS datasheet (Zizhen Yao, Hanqing Liu, Fangming Xie, Stephan Fischer, *et al*. 2021)[3]. The visualization and analysis were conducted using NeMO analytics, available at https://nemoanalytics.org/.

**f**, Representative images (left) and quantification (right) of crossing DRG axons in the presence of function-blocking monoclonal antibodies targeting integrin-β1 (anti-β1 Ab) conditions ($n$ = 12 per group). *n.s.*, not significant. Kruskal-Wallis test followed by post-hoc Dunn correction.

**g**, Representative images, and quantification of crossing DRG axons in the presence of DDR1 inhibitor (nilotinib, merestinib, and sitravatinib) ($n$ = 16 per group). *n.s.*, not significant. Kruskal-Wallis test followed by post-hoc Dunn correction.

**h**, **i**, **j**, Feature of GPR56 (*Adgrg1*) repression in DRG neurons (**h**), neurons in spinal cord, and primary motor cortex neurons (**i**, **j**).

**k**, Scale model of extracellular regions of mouse GPR56 (UniProt: Q8K209, residues S27-S392) based on the crystal structure (PDB: 5KVM)[4].

**I**, Quantification of crossing DRG axons under rΔPLL-hFc (recombinant rat PLL (A26-N175) fused with a human Fc tag at the C-terminal) protein and DHM inhibitor conditions ($n$ = 16 per group). *n.s.*, not significant. Mann Whitney test. Data are min to max, show all points. Scale bars, 200 μm (**f**, **g**, **l**).

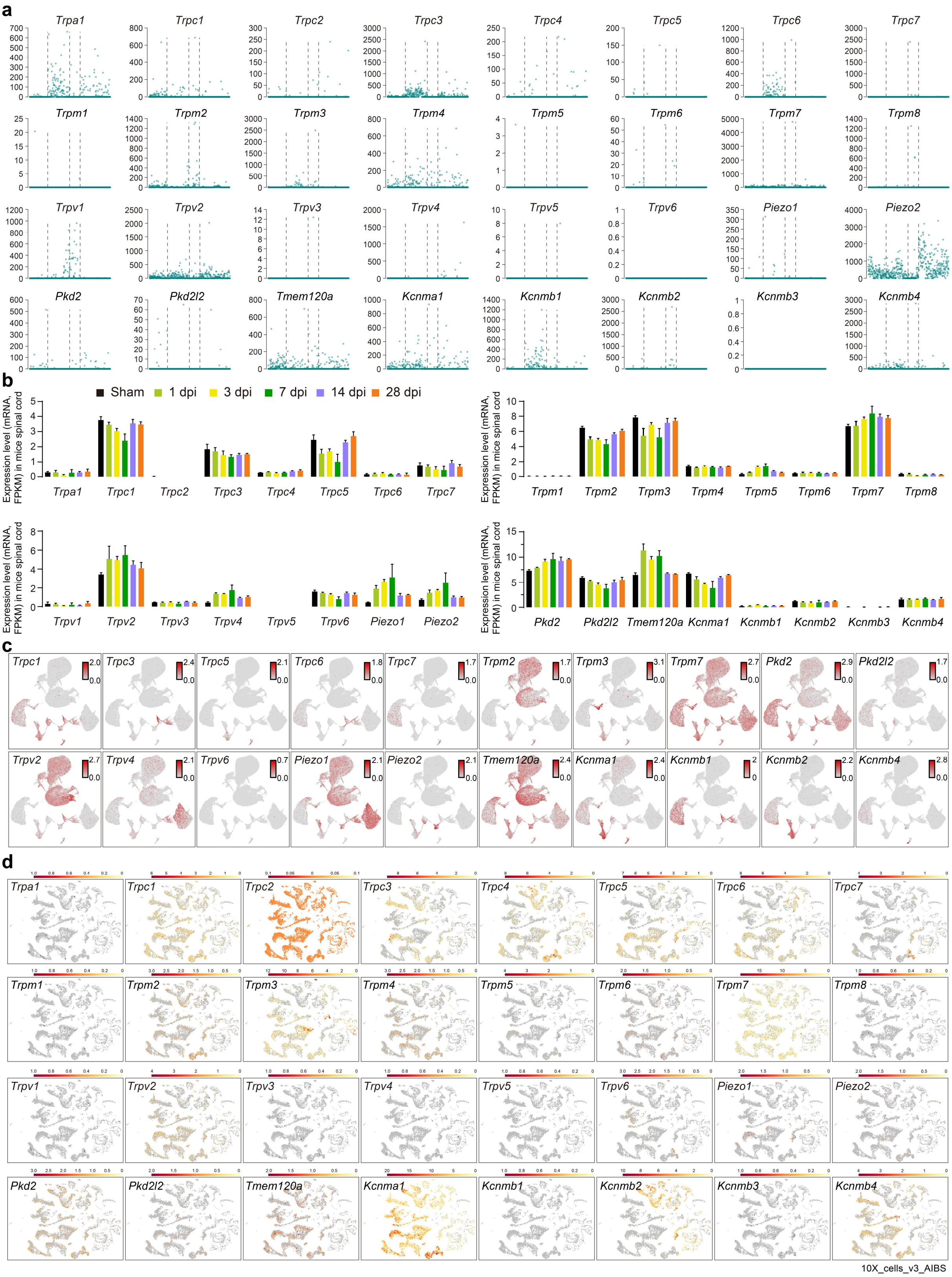

**Extended Data Fig. 9 | Expression of related mechanosensitive channels.**

**a**, **b**, **c**, **d** Potential mechanosensitive channel expression on DRG neurons (**a**), neurons in spinal cord (**b**, **c**), and primary motor cortex neurons (**d**).

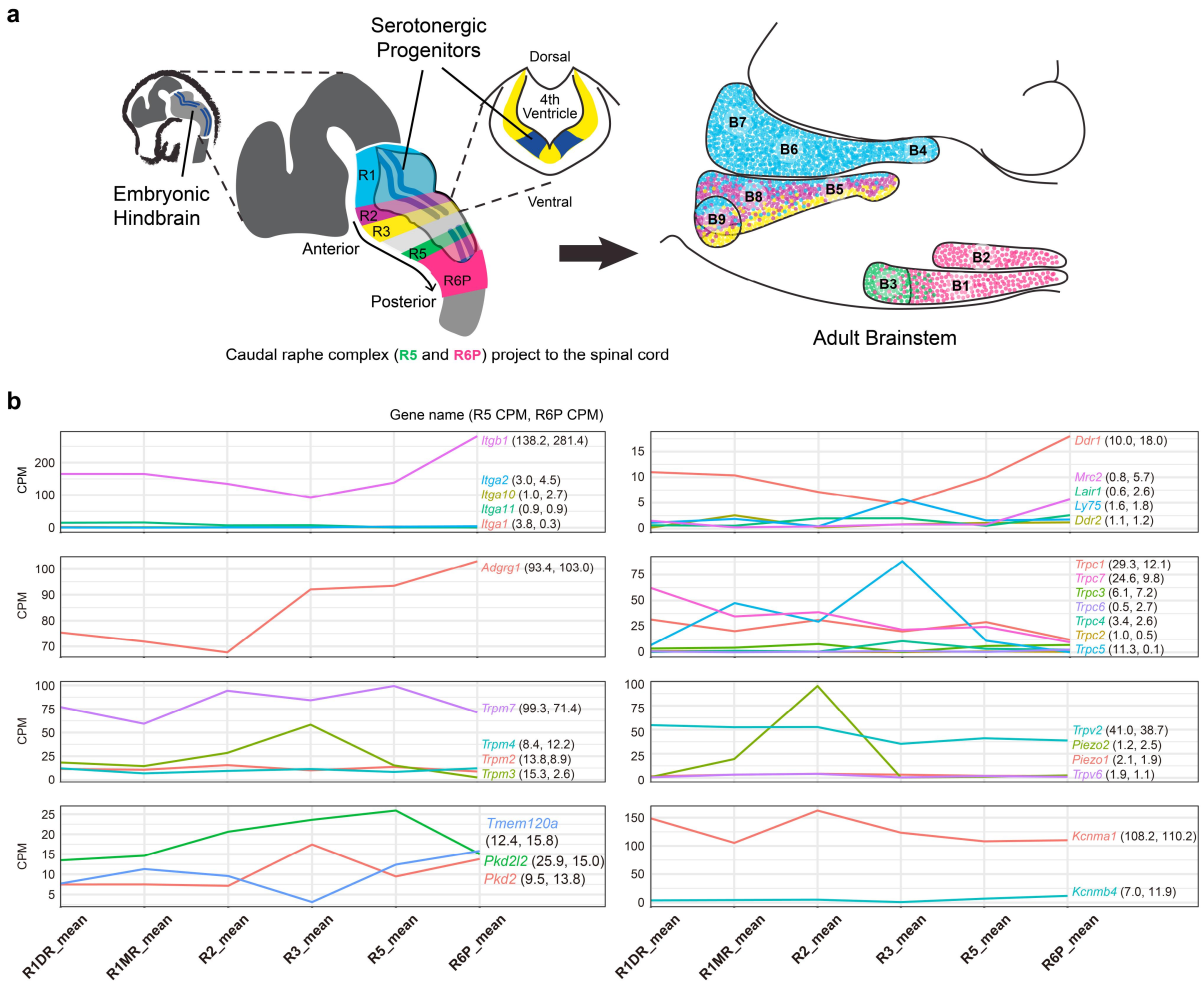


**Extended Data Fig. 10 | An analysis of collagen-associated receptors and potential mechanosensitive channel expression on serotonin neurons within the raphe nucleus**.

**a**, Schematic diagram illustrating the cell source for single-cell sequencing (adapted from Benjamin W. Okaty, Morgan E. Freret, *et al*. 2015)[5]. On the left is a schematic of the mouse embryonic hindbrain, with the serotonergic progenitors marked in dark blue. The rhombomeric areas giving rise to 5-HT neurons, including R1, R2, R3, R5, and R6P, are shaded in different colors. R5 and R6P represent caudal raphe complex projecting into the spinal cord. On the right, the mature anatomical distribution of these neurons is presented in a sagittal brainstem schematic labeled B1-B9 nuclei. Single cell derived from mice aged P25-P32[5].

**b**, Gene expression of indicated receptor in different raphe nucleus domains. Raw data derived from the Neuron paper (doi.org/10.1016/j.neuron.2015.10.007, Table S3, contributed by Benjamin W. Okaty, *et al*). CPM (Counts per million).

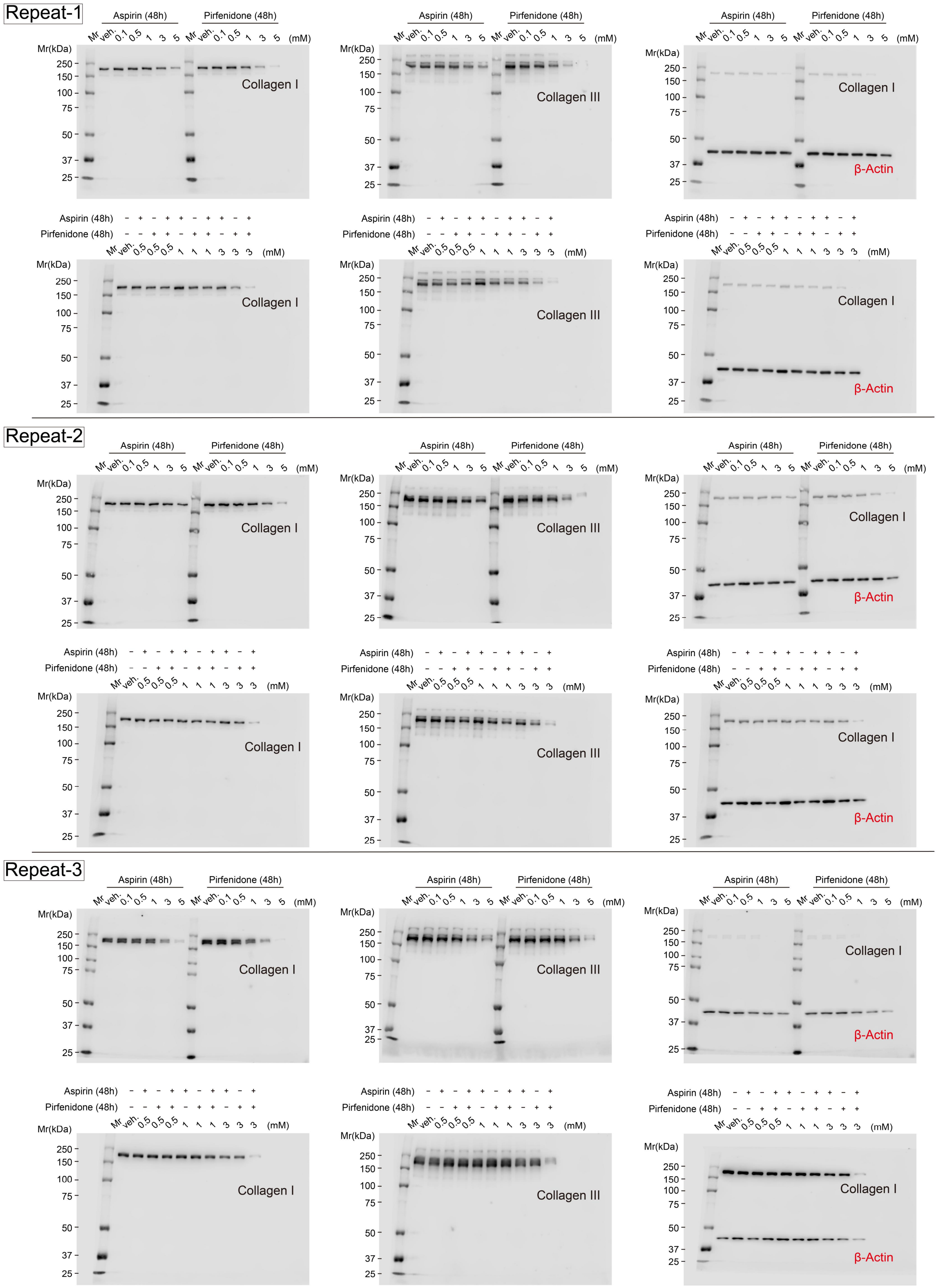
Repeat-1
Repeat-2
Repeat-3
Aspirin (48h)
Pirfenidone (48h)
Mr veh. 0.1 0.5 1 3 5 (mM)
Mr veh. 0.5 0.5 0.5 1 1 1 3 3 3 (mM)
Mr(kDa)
250
150
100
75
50
37
25
Collagen I
Collagen III
β-Actin

**Extended Data Fig. 11 | Uncropped images of immunoblot.**
β-Actin antibody was incubated on the same membrane after collagen I exposure.

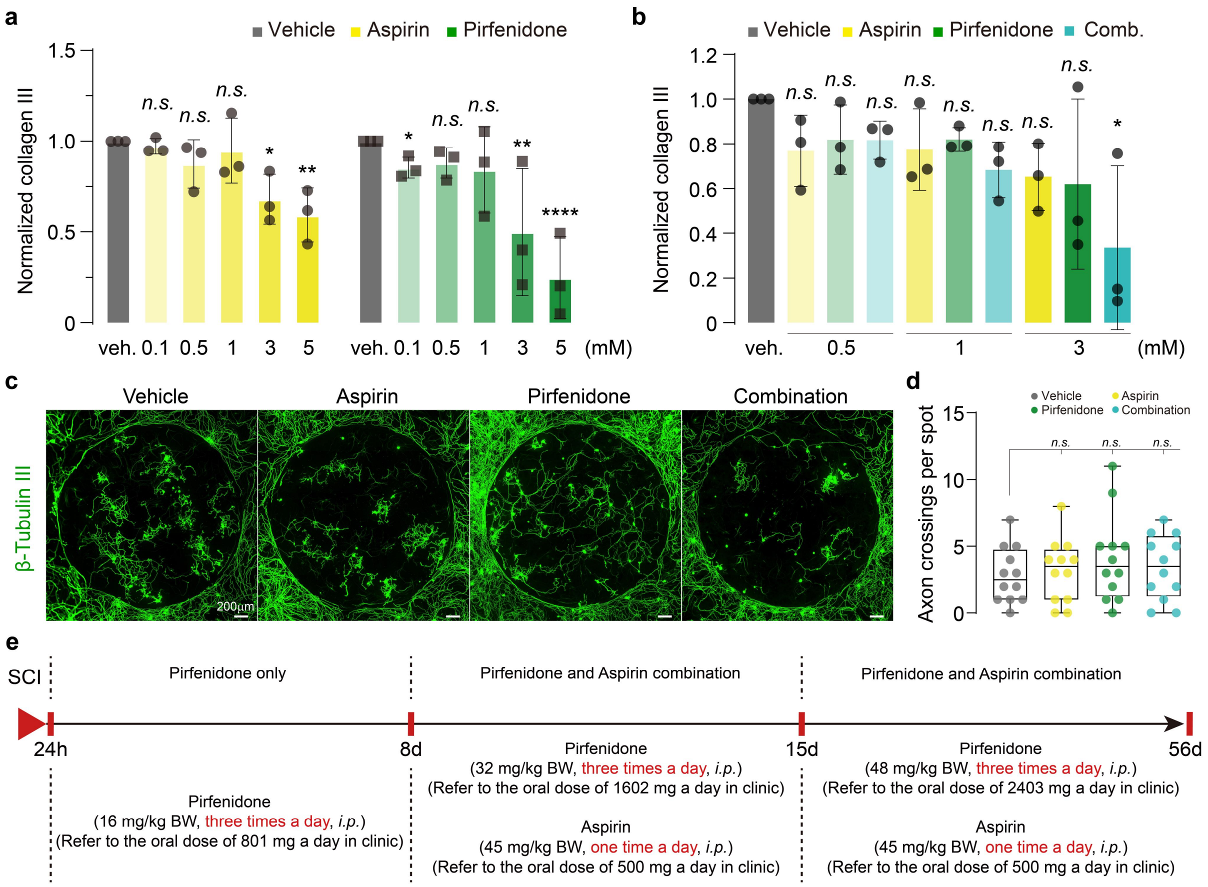


**Extended Data Fig. 12 | Fibroblasts and DRG neuron responses to aspirin and pirfenidone, and timeline for *in vivo* experiments**.

**a**, Quantification of the collagen III intensity (normalized to each β-actin and the vehicle control) in Fig. 3c. Data are mean ± s.d. $n = 3$, three independent tests. *$p = 0.0394$, **$p = 0.0073$, *n.s.*, not significant. Two-way RM ANOVA followed by post-hoc Dunnett correction.

**b**, Quantification of the collagen III intensity (normalized to each β-actin and the vehicle control) in Fig. 3e. Data are mean ± s.d. $n = 3$, three independent tests. *$p = 0.017$, *n.s.*, not significant. Kruskal-Wallis test followed by post-hoc Dunn correction.

**c**, Representative images of DRGs growing on collagen I gradient spots treated with aspirin (3 mM) and pirfenidone (3 mM).

**d**, Quantification of numbers for DRG axons crossing the inhibitory rim ($n = 12$ per group) in **c**. *n.s.*, not significant. one-way ANOVA followed by post-hoc Bonferroni correction. Data are min to max, show all points. **e**, Experimental design and timeline for *in vivo* experiment.

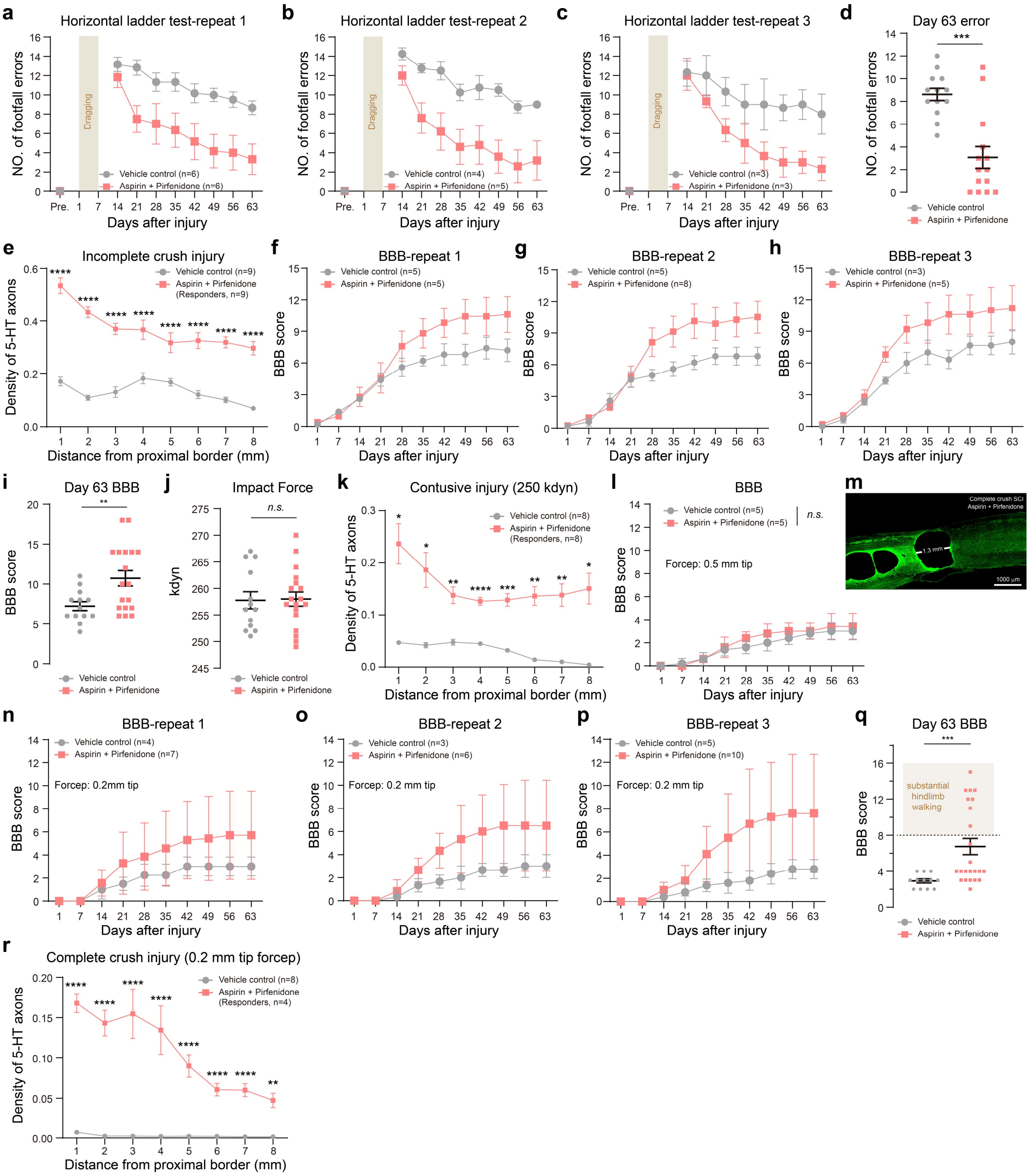

a
Horizontal ladder test-repeat 1
NO. of footfall errors
Dragging
Vehicle control (n=6)
Aspirin + Pirfenidone (n=6)
Days after injury
b
Horizontal ladder test-repeat 2
Vehicle control (n=4)
Aspirin + Pirfenidone (n=5)
c
Horizontal ladder test-repeat 3
Vehicle control (n=3)
Aspirin + Pirfenidone (n=3)
d
Day 63 error
Vehicle control
Aspirin + Pirfenidone
e
Incomplete crush injury
Density of 5-HT axons
Vehicle control (n=9)
Aspirin + Pirfenidone (Responders, n=9)
Distance from proximal border (mm)
f
BBB-repeat 1
BBB score
Vehicle control (n=5)
Aspirin + Pirfenidone (n=5)
g
BBB-repeat 2
Vehicle control (n=5)
Aspirin + Pirfenidone (n=8)
h
BBB-repeat 3
Vehicle control (n=3)
Aspirin + Pirfenidone (n=5)
i
Day 63 BBB
j
Impact Force
kdyn
n.s.
k
Contusive injury (250 kdyn)
Vehicle control (n=8)
Aspirin + Pirfenidone (Responders, n=8)
l
BBB
Vehicle control (n=5)
Aspirin + Pirfenidone (n=5)
Forcep: 0.5 mm tip
m
Complete crush SCI
Aspirin + Pirfenidone
1.3 mm
1000 μm
n
BBB-repeat 1
Vehicle control (n=4)
Aspirin + Pirfenidone (n=7)
Forcep: 0.2mm tip
o
BBB-repeat 2
Vehicle control (n=3)
Aspirin + Pirfenidone (n=6)
Forcep: 0.2 mm tip
p
BBB-repeat 3
Vehicle control (n=5)
Aspirin + Pirfenidone (n=10)
Forcep: 0.2 mm tip
q
Day 63 BBB
substantial hindlimb walking
r
Complete crush injury (0.2 mm tip forcep)
Vehicle control (n=8)
Aspirin + Pirfenidone (Responders, n=4)

**Extended Data Fig. 13 | Detailed behavioral data and analyses**.

**a-c**, The individual data show three repeat experiments of hindlimb errors in the horizontal ladder test after aspirin and pirfenidone treatment (Incomplete crush SCI). Data are mean ± s.e.m.

**d**, Quantification of hindlimb errors at 63 dpi ($n$ = 13 vehicle control, $n$ = 14 combination). ***$p$ = 0.0005, Mann Whitney test. Data are mean ± s.e.m.

**e**, Quantification of the density of 5-HT axons in the spinal cord distal to the injury site (normalized to the density of 1 x 1 $mm^2$ proximal to the injury border) at 9 weeks post incomplete crush injury. ($n$ = 9). ****$p$ < 0.0001, two-way RM ANOVA followed by post-hoc Bonferroni correction. Data are mean ± s.e.m.

**f-h**, The individual three repeats of BBB scores after aspirin and pirfenidone treatment in contusive SCI (250 kdyn). Data are mean ± s.e.m.

**i**, Quantification of BBB score at 63 dpi ($n$ = 13 vehicle control, $n$ = 18 combination). **$p$ = 0.0043, unpaired t test with Welch's correction. Data are mean ± s.e.m.

**j**, The force of all rats that received an infinite 250 kdyn horizon contusive injury (variability less than 10%). Mean of vehicle control = 257.8 kdyn, Mean of combination = 258.0 kdyn. *n.s.*, not significant, unpaired t test. Data are mean ± s.e.m.

**k**, Quantification of the density of 5-HT axons in the spinal cord distal to the injury site (normalized to the density of 1 x 1 $mm^2$ proximal to the injury border) at 9 weeks post 250 kdyn contusive injury. ($n$ = 9). *$p$ < 0.05, **$p$ < 0.01, ***$p$ < 0.001, ****$p$ < 0.0001, two-way RM ANOVA followed by post-hoc Bonferroni correction. Data are mean ± s.e.m.

**l**, Locomotor recovery (BBB score) in complete crush SCI (0.5 mm tip forceps) ($n$ = 5). *n.s.*, not significant, two-way RM ANOVA followed by post-hoc Bonferroni correction Data. are mean ± s.d.

**m**, Representative image of extensive cavity formation at T10 in an aspirin and pirfenidone treatment group at 9 weeks post complete crush SCI (0.5 mm tip forceps).

**n-p**, The individual three repeats of BBB scores after aspirin and pirfenidone treatment in complete crush SCI (modified forceps with 0.2 mm tip). Data are mean ± s.d.

**q**. Quantification of BBB scores at 63 dpi ($n$ = 12 vehicle control, $n$ = 23 combination). ***$p$ = 0.001, Mann Whitney test.

**r**, Quantification of the density of 5-HT axons in the spinal cord distal to the injury site (normalized to the density of 1 x 1 $mm^2$ proximal to the injury border) at 9 weeks post complete crush injury. ($n$ = 8 vehicle control, $n$ = 4 combination). **$p$ = 0.0024, ****$p$ < 0.0001, two-way RM ANOVA followed by post-hoc Bonferroni correction. Data are mean ± s.e.m. Scale bars, 1000 μm (**m**).

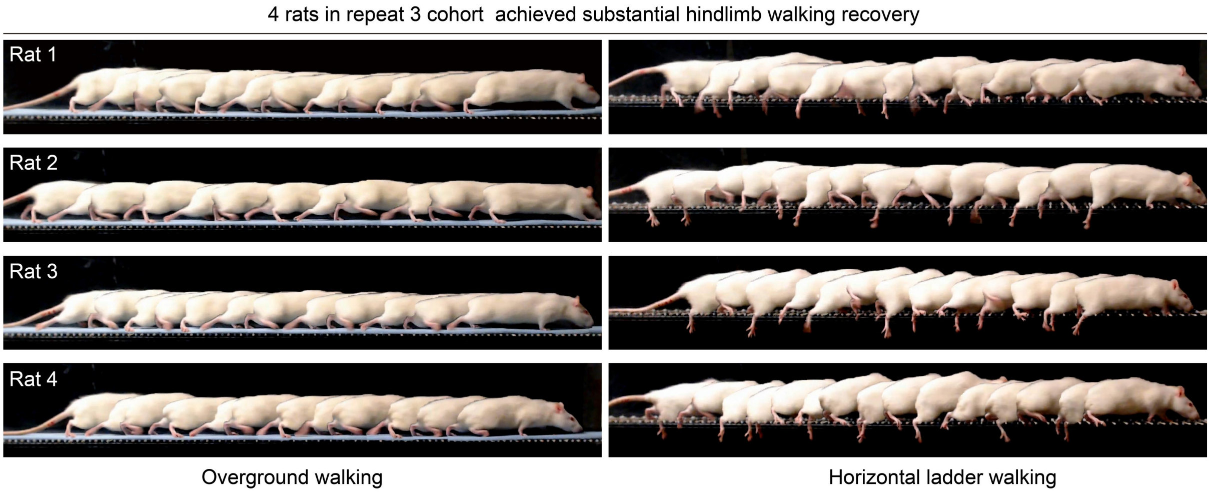


**Extended Data Fig. 14 | Restoration of walking following aspirin and pirfenidone combination treatment in repeated cohorts of rats**.

Locomotor performance of 4 responders in cohort 3 at 9 weeks post complete crush SCI (modified forceps with 0.2 mm tip). Chronophotography of overground walking showing substantial hindlimb movements, and chronophotography of horizontal ladder walking showing a lack of coordination in these steps.

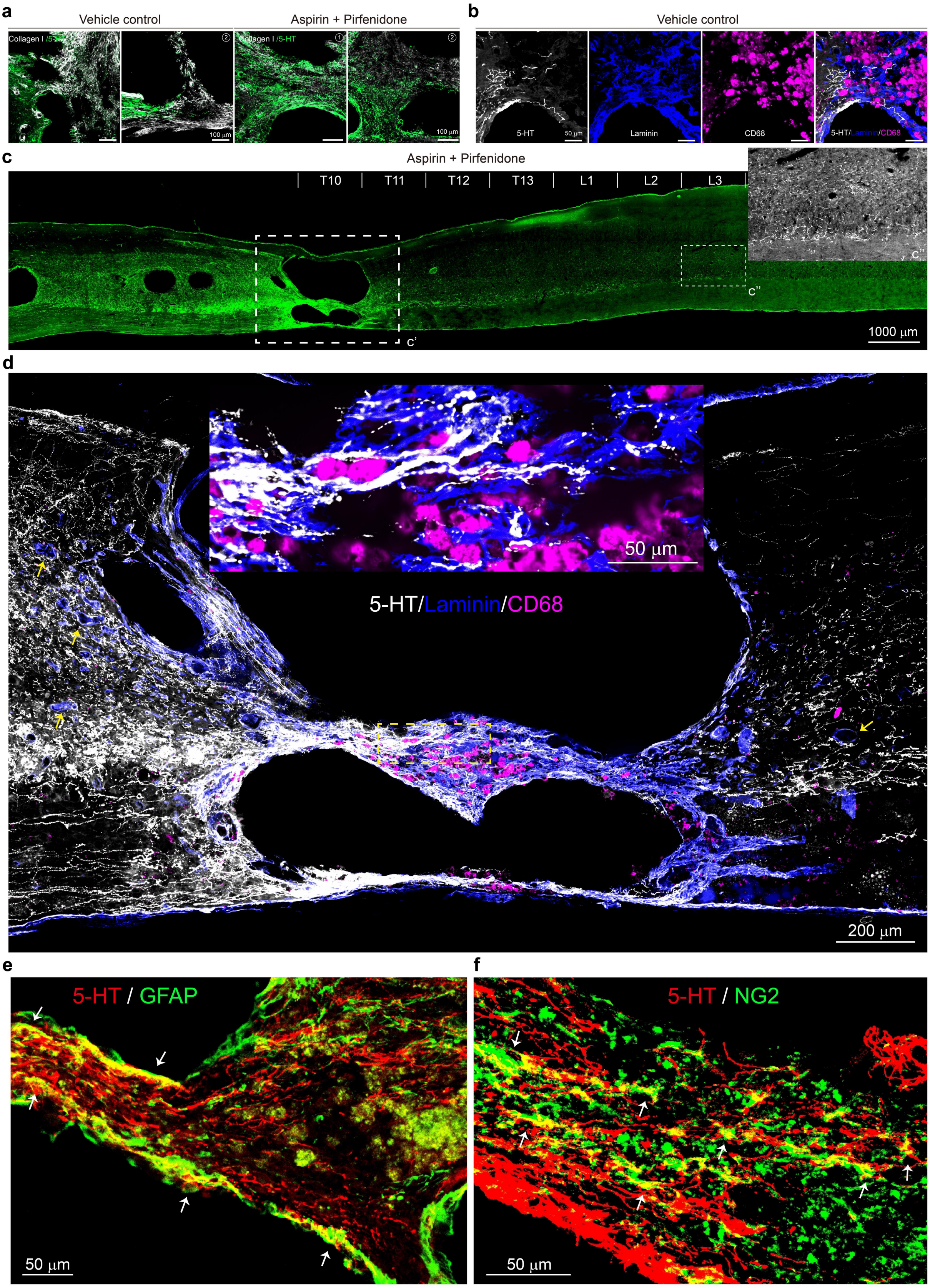
a
Vehicle control
Aspirin + Pirfenidone
Collagen I /5-HT
Collagen I /5-HT
100 μm
100 μm
b
Vehicle control
5-HT
50 μm
Laminin
CD68
5-HT/Laminin/CD68
c
Aspirin + Pirfenidone
T10
T11
T12
T13
L1
L2
L3
c'
c''
1000 μm
d
50 μm
5-HT/Laminin/CD68
200 μm
e
5-HT / GFAP
50 μm
f
5-HT / NG2
50 μm

**Extended Data Fig. 15 | Regenerated 5-HT axons regrow through astrocyte and NG2$^+$ cell bridges laden with laminin. Axons avoid CD68$^+$ microglia/macrophages**.

**a**, Surveys of 5-HT axons and collagen I in the lesion core at 9 weeks post-SCI**.**

**b**, 5-HT axons stop at the proximal cell bridge where it is filled with laminin and CD68$^+$ microglia/macrophages but also collagen I in vehicle-treated rats**.**

**c**, Panoramic sagittal view of 5-HT axon projections in the spinal cord of a rat with functional recovery. The lesion core c' and further caudally at the lumbar level c'' are enlarged. T, thoracic; L, lumbar.

**d**, 5-HT axon regrowth in the lesion core occurs after a decrease in collagen I/III along laminin$^+$ cells but not on CD68$^+$ microglia/macrophages. The arrows indicate that the newly formed microvasculature in the glial/fibrotic scar penumbra is closely associated with dramatic increases laminin expression, which suggests that newly formed laminin may originate from pericytes.

**e**, **f**, 5-HT axon regrowth through astrocytes in the proximal borders and NG2$^+$ cells in the lesion core. Scale bars, 100 µm (**a**), 50 µm (**b**), 1000 µm (**c**), 50 µm and 200 µm (**d**), 50 µm (**e,f**).
